# Plectin affects cell viscoelasticity at small and large deformations

**DOI:** 10.1101/2025.05.28.656588

**Authors:** James P. Conboy, Mathilde G. Lettinga, Nicole van Vliet, Lilli Winter, Gerhard Wiche, Frederick C. MacKintosh, Gijsje H. Koenderink

**Author notes:** These authors contributed equally to this work (James P. Conboy, Mathilde G. Lettinga). Center for Molecular and Cellular Bioengineering, BIOTEC, TU Dresden, Germany.

## Abstract

Plectin is a giant protein of the plakin family that crosslinks the cytoskeleton of mammalian cells. It is expressed in virtually all tissues and its dysfunction is associated with various diseases such as skin blistering. There is evidence that plectin regulates the mechanical integrity of the cytoskeleton in diverse cell and tissue types. However, it is unknown how plectin modulates the mechanical response of cells depending on the frequency and amplitude of mechanical loading. Here we demonstrate the role of plectin in the viscoelastic properties of fibroblasts at small and large deformations by quantitative single-cell compression measurements. To identify the importance of plectin, we compared the mechanical properties of wild type (*Plec*^+/+^) fibroblasts and plectin knockout (*Plec*^−/−^) fibroblasts. We show that plectin knockout cells are nearly 2-fold softer than wild type cells, but their strain-stiffening behaviour is similar. Plectin deficiency also caused faster viscoelastic stress relaxation at long times. Fluorescence recovery after photobleaching experiments indicated that this was due to 3-fold faster actin turnover. Short-time poroelastic relaxation was also faster in *Plec*^−/−^ cells as compared to *Plec*^+/+^ cells, suggesting a more sparse cytoskeletal network. Confocal imaging indicated that this was due to a marked change in the architecture of the vimentin network, from a fine meshwork in wild type cells to a bundled network in the plectin knockout cells. Our findings therefore indicate that plectin is an important regulator of the organization and viscoelastic properties of the cytoskeleton in fibroblasts. Our findings emphasize that mechanical integration of the different cytoskeletal networks present in cells is important for regulating the versatile mechanical properties of cells.

**SIGNIFICANCE:** Mammalian cells combine superior mechanical strength with the ability to actively deform themselves. They owe this paradoxical mechanical behaviour to their cytoskeleton, an intracellular web of protein filaments that includes actin filaments and intermediate filaments. It is known that both cytoskeletal filament types contribute to cell stiffness on their own, but the impact of their mechanical integration via cytoskeletal crosslinker proteins remains unknown. Here we test the effect of crosslinking of actin and vimentin intermediate filaments by the crosslinker protein plectin in fibroblasts by single-cell compression measurements. By comparing normal cells and cells in which plectin is knocked out, we find that plectin significantly increases cell stiffness and provides a protective mechanism against actin network disruption by compressive loading.

## INTRODUCTION

Cells possess an extraordinary capacity to preserve their structural integrity in physically stressful environments. In mammalian cells, the main structure governing mechanical protection is the cytoskeleton, a biopolymer network that extends throughout the cell’s interior (1). The cytoskeleton is composed of four distinct biopolymers: actin, microtubules, intermediate filaments and septins, each playing specific roles in different cellular functions (2). The filaments differ in their mechanical and dynamical properties. With respect to mechanics, intermediate filaments and septins are the softest, with a persistence length of only ∼1 µm (3, 4), while actin filaments and microtubules are stiffer, with persistence lengths of ∼10 µm and several millimeters, respectively (5). With respect to dynamics, actin filaments and microtubules have high polymerization and depolymerization rates, while intermediate filaments and septins form more stable polymers (2). Each cytoskeletal filament is organized into higher-order structures such as networks or bundles by their own dedicated set of accessory proteins such as crosslinker proteins and polymerization factors (6–9). Cell-free reconstitution experiments and theoretical models have shown that crosslinker proteins sensitively control the elastic properties of cytoskeletal networks, including the elastic modulus, rupture strength, and ability to strain-stiffen (10). In addition, the dynamic binding and unbinding of actin crosslinkers introduces time-dependent viscoelastic properties (11–13).

The mechanical functions of the different cytoskeletal filament systems have traditionally been studied separately. Recently, however, there has been growing appreciation for the importance of physical and biochemical coupling of the filaments in the ability of cells to resist, transmit and generate mechanical forces (2). Spectraplakin proteins play a particularly important role in mechanical integration of the cytoskeleton by forming crosslinks between different filament types (14, 15). Spektraplakins are large (>500 kDa) and evolutionarily conserved crosslinker proteins that are part of the spectrin superfamily. Plectin is a widely expressed member of this protein family that serves as a crosslinker between actin and intermediate filaments (16, 17). Human plectin is encoded by a single gene located on chromosome 8 (18), but alternative splicing creates 12 different plectin isoforms expressed in different combinations in different cell and tissue types (19).

All plectin isoforms have a C-terminal intermediate filament-binding domain and an actin-binding domain (ABD) located close to their N termini. Moreover, they all share a spectrin repeat-containing plakin domain and a central alpha-helical rod domain that mediates plectin dimerization. By contrast, the N-terminal head domain is variable. This domain controls the cellular localization of each plectin isoform by docking to distinct interaction partners such as the nucleus (plectin isoform P1), junctional complexes like focal adhesions (isoform P1f), or microtubules (isoform P1c) (19). As a consequence, plectins simultaneously crosslink intermediate filaments to actin filaments and connect these cytoskeletal networks to microtubules, the nucleus, and adhesion complexes (20). Through its connector role, plectin is essential for a stable cytoarchitecture and mechanical integrity of epithelial and endothelial tissues (21–24). Accordingly, plectin mutations result in multiple multisystemic diseases known as plectinopathies (25–27). A well-known example is epidermolysis bullosa simplex (EBS-MD), a disease characterised by spontaneous skin blistering, impaired wound healing and muscular dystrophy (28). At the cell level, dermal fibroblasts from EBS-MD patients exhibit an anomalous vimentin network architecture, with vimentin intermediate filaments organized in bundles instead of meshworks (28). Plectin knockout in keratinocytes results in similar changes in the keratin cytoskeleton, from keratin meshworks to bundles (29).

In mesenchymal cells, plectin crosslinks vimentin to different actin structures, including the actin cortex of mitotic cells (30, 31), invadopodial actin networks of invading cancer cells (32), and actin stress fibres in adherent cells (33, 34). One function of plectin-mediated crosslinking of actin and vimentin is to control the transmission of actin-myosin-based forces in cells, both in 2D-adherent conditions and in 3D matrices (34, 35). Given this mechanical function, plectin-mediated crosslinking would also be expected to influence the stiffness of the cytoskeleton. In biochemically reconstituted actin-vimentin networks, plectin-mediated crosslinking indeed causes network stiffening (36). Surprisingly, however, measurements of plectin’s contribution to cellular elasticity are contradictory. In mouse skin fibroblasts probed with magnetic twisting cytometry, plectin was found to increase cell stiffness (35). Similar behaviour was observed for myoblasts probed with magnetic tweezers (37). However, this same study showed an opposite behaviour for keratinocytes, where plectin knockout made cells stiffer. Atomic force microscopy (AFM) on malignant epithelial cells showed a negligible affect of plectin knock-down on cell stiffness (38). These discrepancies could point to cell-type-specific functions of plectin, caused, for instance, by different expression levels of the various plectin isoforms. Differences in the experimental assays could also contribute to the reported discrepancies. It is well-known that values of cell stiffness and viscosity vary substantially depending on the level of applied mechanical stress, the rate of deformation, the geometry of the measurement probe, the location probed in the cell, and the extracellular microenvironment (39). Prior studies of the role of plectin in cell mechanics were based on local probing by AFM-based nanoindentation or magnetic tweezer manipulation of micron-sized beads attached to focal adhesions. This type of local probing has been shown to be more sensitive to cortical actin than to cytoplasmic intermediate filaments (40, 41). Furthermore, these methods are limited to small deformations, whereas cells in the body tend to experience large deformations where nonlinear elastic properties become important.

Here we investigate the mechanical role of plectin in fibroblasts at small and large deformations by single-cell compression with our newly developed device that affords high precision in strain rate and amplitude (42). To identify the role of plectin, we compare the linear viscoelastic properties of wild type versus plectin knockout cells in response to small amplitude oscillatory and step strains. We find that plectin has a significant impact on bulk cell stiffness and also influences the time-dependent poroelastic and viscoelastic behaviour of the cells. Strain ramp experiments compressing the cells into the nonlinear elastic regime confirm that the plectin knockout cells are softer than wild type cells at small strains, but show similar strain-stiffening behavior. Finally, under cyclical straining, we observed that wild type cells are initially stiffer than plectin knockout cells, but attain a similar stiffness as plectin knockout cells upon repeated loading, suggesting that plectin shields the cytoskeleton from damage by providing transient crosslinks that dissociate under load.

## MATERIALS AND METHODS

### Cell culture and sample preparation for compression experiments

All experiments were performed with immortalized mouse fibroblast cell lines derived from transgenic plectin wild-type (*Plec*^+/+^) mice or plectin-deficient (*Plec*^−/−^) mice in which the plectin gene had been targeted and inactivated by homologous recombination; consequently, *Plec*^−/−^ fibroblasts used in this study were plectin-null, i.e. lacking all isoforms of plectin. Generation of transgenic mouse lines and isolation of immortalized cell lines have previously been described in detail (see (43–45)). Cells were cultured in Dulbecco’s modified Eagle’s medium with glutamax (DMEM, 10565018 Gibco) supplemented with 10% fetal bovine serum (FBS, 10270106 Gibco) and 5% penicillin-streptomycin (Pen-Strep antibiotic, 15070063 Gibco). Cells were maintained by subculturing twice a week and were screened for mycoplasma contamination every four months. Experiments were performed with cells up to passage number 20.

One day before experiments, cells were detached with 0.25% Trypsin-EDTA (25200056 Thermo Fisher Scientific) and counted using a Countess cell counter (Thermo Fisher Scientific). Next, 10,000 cells were transferred to plastic cell culture 6-well plates (83.3920.005 Sarstedt) containing culture medium. The cells were left to grow to ∼ 70% confluence. On the day of the experiments, the cells were stained by replacing the medium with medium containing CellTracker Orange (10082742 Thermo Fisher Scientific, 1:1,000 dilution). After 30 min incubation, the medium was removed and the cells were detached from the plate by incubating with 200 µl Trypsin-EDTA (0.25%) for 3 minutes. The cells were then suspended in 4 mL

CO_2_-independent medium (18045088 Gibco), prewarmed to 37 °C. Next, the cells were seeded into 35 mm home-made dishes. Dishes were made by glueing 19 mm glass coverslips (12323138 Thermo Fisher Scientific) precleaned by sonication in 2-propanol (5 min) to Petri dishes (Sarstedt) with Norland Optical Adhesive glue. A drop of glue was placed in the dish, the coverslip was carefully placed on top to make sure the glue was evenly spread, and the glue was cured for 5 min using a UV ozone ProCleaner (Bioforce Nanosciences). The dishes were coated with Pluronic F-127 (P2443 Sigma) dissolved (1:100 w/v) in PBS (10010023 Gibco) for 15 min before the experiment.

### Single-cell compression experiments

Cell compression experiments were performed using a Chiaro Nanoindenter operated by the Piuma software (Optics11 Life) mounted on a Leica Thunder Imager wide field microscope. The cells were imaged *in situ* using a 200 mW solid state LED5 light source (Leica), a monochrome sCMOS camera (Leica), and a 10x dry objective (HC PL APO 0.45NA, Leica). We used custom-made wedged cantilevers prepared by modification of tipless cantilevers with a nominal spring constant of 0.018 N/m (Optics11 Life). To correct for the 3^°^tilt of the cantilevers (46), we fabricated a wedge using Norland optical adhesive (NOA81 Norland), as described in our previous work (42). We calibrated the cantilevers before and after wedge manufacture by the stiff-surface contact method (as described by the manufacturer) to verify that the spring constant was unchanged.

In preparation for compression experiments, a cell dish containing 4 mL of pre-warmed CO_2_-independent medium was placed on the microscope stage. Before lowering the cantilever probe into the medium, we added a drop of medium onto the probe to prevent air bubbles. The probe was first calibrated by the stiff-surface contact method which involves pressing against the coverslip. Next, the cantilever was moved up and 200 µl of cell suspension was added to the dish. After allowing cells to sediment for ∼ 10 minutes, the cantilever was moved down to a distance of ~200 µm above the coverslip. We then positioned a single cell underneath the cantilever with help of the fluorescent signal of the CellTracker dye in the cytosol. We carefully lowered the cantilever until reaching contact with the cell, apparent from the optically detected cantilever deflection. In all experiments, we controlled the indentation (Indentation mode in the Piuma software), which allows precise control of the rate of deformation. During measurements, the elapsed time, piezo displacement, and cantilever deflection were recorded at a time resolution of 1000 Hz. We verified by live-dead staining that the cells remained viable over the entire duration (at most 120 min) of the experiments (42).

We used three different protocols to deform the cells. First, for small amplitude oscillatory measurements, we lowered the cantilever probe by 1 µm after finding the cell surface, corresponding to a compressive strain of ∼ 0.05, which is well within the linear elastic regime. Next, a sinusoidal displacement with an amplitude of 200 nm was applied at five logarithmically spaced frequencies between 0.1 Hz and 10 Hz, corresponding to strain rates between 0.02 µm/s and 2 µm/s. Second, for stress relaxation measurement, we applied a small step strain by quickly compressing the cell by a distance of 1 µm (corresponding to a strain of ∼ 0.05) at a large strain rate of 10 µm/s. We subsequently measured the time-dependent relaxation of the stress over a period of 80 s. Third, to capture the nonlinear response of the cells to high levels of compressive strain, we performed strain ramp measurements at a constant strain rate of 1 µm/s up to a maximal deformation of 8 µm, corresponding to a strain of ∼ 0.4.

### Analysis of single-cell compression data

All data analysis was performed using custom-written Python code available upon request. The load measured by the Nanoindenter was converted to compressive engineering stress based on the initial contact area between the cell and the cantilever. The contact area was determined from automated analysis of epifluorescence images of the cytosol by thresholding the images using Otsu’s method in the Python library scikit-image (47) and multiplying the number of pixels by the pixel size in the thresholded image. The uniaxial strain was calculated by approximating cells as a sphere and dividing the amount of compression by the cell diameter. We note that the measurements and all analyses were previously benchmarked using cell-sized polyacrylamide microgel particles of known stiffness (42). Curve fitting was done using the curve fit function from Python’s SciPy library for oscillatory and strain ramp measurements (48) and the Julia RHEOS package (49) for stress relaxation experiments.

### Small amplitude oscillatory measurements

We calculated the complex viscoelastic modulus, *E*^*\**^ (*ω*) = *FT* (*σ* ((*t*)/ *FT*(*ϵ*(*t*)), at each oscillation frequency *ω* = 2*π f*, from the Fourier transformed (FT) stress, *σ* (*ω*), and strain, *ϵ* (*ω*). The Fourier transformations were performed using the fast Fourier transform (FFT) function in Python’s numpy library. We separated the complex modulus, defined as *E*^*\**^ = *E*^′^+ *iE*^′′^, into its real and imaginary parts to obtain the compressive storage modulus, *E*^′^ (*ω*), and loss modulus, *E*^′′^ *ω*. *E*^′^ (*ω*) corresponds to the elastic modulus of the cell (i.e., the energy stored per cycle of oscillation), whereas *E*^′′^ (*ω*) corresponds to the viscous component (i.e., the energy dissipated per cycle of oscillation).

To mechanically fingerprint our cells, we first tried to fit the frequency spectra to the simplest possible phenomenological model referred to as the structural damping model (50–52) (for more details see Supplementary Methods). Briefly, this three parameter model is described by:

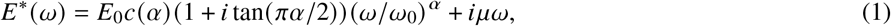

where *E*_0_ is a scaling factor for the elastic and viscous moduli at a frequency scale factor *ω*_0_ (which we arbitrarily set to 1), *c α* = (Γ 1 −*α*) cos (*πα* /2) with Γ the gamma function, *α* is the power law exponent and *μ* is the Newtonian viscous term. As this model failed to capture the rheology of our cells, we applied a fractional rheological model adapted from recent work of Bonfanti et al. (52, 53):

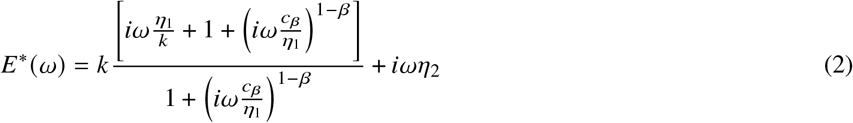

This model has five fitting parameters: *k* is the elastic modulus representing the stiffness of the cell, *η*_1_ is a dashpot viscosity capturing long-term viscous dissipation, *c*_*β*_ is a spring-pot parameter describing the intermediate viscoelastic behaviour, *β* is a fractional exponent that characterises power-law scaling with frequency, and *η*_2_ is a Newtonian viscous term capturing the response of the cytoplasm. We further decomposed Equation 2 into real and imaginary parts:

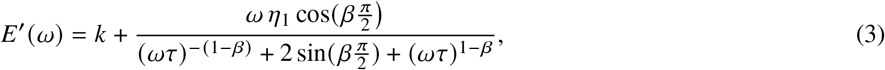

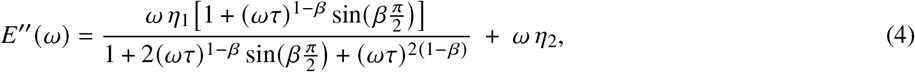

For both models, we simultaneously fitted the measured storage and loss moduli to these equations to find a self-consistent description of the elastic and viscous response.

### Step strain measurements

We found that stress relaxation after a step strain exhibited two distinct time regimes characterized by different functional forms. In the first regime, for times between 10^−3^-10^−1^s, we fitted the stress response to a poroelastic model, where stress decays exponentially with time (54):

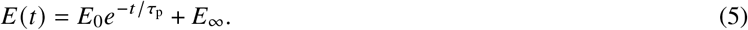

Here, *E*_0_ denotes the instantaneous modulus immediately after loading, *E*_*∞*_ is the equilibrium modulus at long times, and *τ*_p_ is the poroelastic relaxation time. In the second regime (times between 0.1-80 s), where the cell behaves as a viscoelastic material, we fitted the same 5-parameter fractional rheological model as with the oscillatory measurements. To obtain the relaxation modulus *E* (*t*), we calculated the Laplace transform of Equation 2, which yields

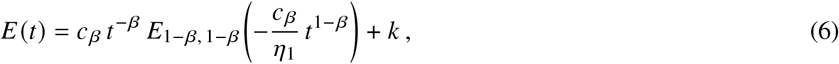

where *E*_*a,b*_ (*z*) denotes the Mittag-Leffler function, a function that naturally appears in the solutions of fractional differential equations (39). To obtain parameter fits, we fit Equation 6 to the stress relaxation data using the Julia RHEOS package (49).

### Strain ramp measurements

The differential cell stiffness *K* was calculated as the derivative of the stress versus strain curves using the Python np.gradient function (55). The linear modulus *K*_0_ and the onset strain (*ϵ* ^*\**^) and onset stress (*σ*^*\**^) where stress-stiffening sets in were determined by performing a piecewise linear fit to the stress/stiffness curve for each individual cell. We fitted a horizontal line and a linear equation of the form *y* = *mx* + *c* with the Python scipy library (48), optimizing the onset point by maximizing the combined *R*^2^ values of both fits. The linear modulus was given by the y-intercept of the horizontal line. To rescale the stress/stiffness curves onto a single master curve, we rescaled *K* by *K*_0_ and *σ* by *σ*^*\**^.

### Confocal microscopy experiments

Confocal imaging of the cells was performed on a Leica Stellaris 8 LSCM microscope, using a 63x, NA 1.30 glycerol immersion objective, white light laser, and HyD S and HyD X detectors in counting mode. Four days prior to experiments, the vimentin cytoskeleton was labelled by transfecting the cells with a GFP-vimentin plasmid (56) using electroporation with the Invitrogen Neon transfection system (Thermo Fisher Scientfic) according to the manufacturer’s instructions. A single pulse of 1350 V was applied for 30 ms. We found that cells required four days to express vimentin at stable levels. For actin labeling, on the day of experiments, cell culture medium was replaced with fresh medium containing SiR actin (Tebu Bio, SC001) (1:1000 from a 1 mM stock solution) along with the pump inhibitor verampamil (Tebu Bio, SC001, part of SiR actin kit) (1:1000 from a 1 mM stock solution). After a 1 hour incubation, the medium was replaced with fresh medium containing Hoechst 33342 (Thermo Fisher Scientific, H3570) (1:10,000) to label the nucleus. The dye incubation was done directly before imaging. To mimic the conditions of the single-cell compression experiments, cells were trapped between two coverslips coated with Pluronic F-127 (Sigma, P2443) spaced apart by 10 µm-sized silica particles (Sigma 904368). We acquired 3D confocal z-stacks of 256 × 256 pixel xy-slices separated by a z-distance of 0.33 µm at a scan speed of 400 Hz.

### Analysis of confocal microscopy data

The orthogonal projection tool in FIJI (57) was used to generate side projections of cells. To generate ‘average cells’, a maximum intensity projection was generated from a 2 µm section centered at the cell equator. Multiple-channel stacks (containing projections of the actin, vimentin and the nucleus) were aligned using the HyperStackReg plugin (58), which registered cells on top of each other, correcting for differences in cell size. In FIJI, we then made average intensity projections to generate an average cell. Line profiles of the actin, vimentin and nucleus signals were determined both for single cells and ‘average cells’. To this end, the radial profile plot tool in FIJI was used to calculate the normalized intensities around concentric circles from the center of the image. Images of at least twelve cells were collected from at least two independent experiments for each condition.

### FRAP measurements of cytoplasmic viscosity and actin network turnover

Fluorescence recovery after photobleaching (FRAP) measurements were performed using the same Leica Stellaris 8 confocal microscope described above, operated in FRAP mode in the LASX software. To ensure effective photobleaching, we used the *zoom-in* and *background to zero* settings as well as the FRAP booster in the Leica LASX software. Cells were again trapped between two coverslips spaced apart by 10 µm-sized beads to mimic the geometry of the compression experiments. For determining cytoplasmic viscosity, the cells were stained with CellTracker Orange (10082742 Thermo Fisher Scientific, 1:1000 dilution in medium) on the day of the experiments. For determining actin turnover, the actin cytoskeleton was labelled by transfecting the cells with an actin-GFP plasmid (59) using electroporation with the Invitrogen Neon transfection system (Thermo Fisher Scientfic). A single pulse of 1350 V was applied for 30 ms.

For both CellTracker and actin FRAP experiments, we imaged a 128×128 pixel confocal xy-slice using the HyD S detector. First, two pre-bleach images were taken to determine the fluorescence intensity before bleaching. Both the CellTracker dye and the actin-GPF were photobleached within a circular region with a radius of 1 µm in the central xy-plane of the cell. This diameter was chosen to ensure that recovery is clearly visible in the images, while minimizing cell damage. To achieve efficient bleaching, we performed two exposures, each lasting 0.23 s, using four laser lines operated at 100% intensity (531 nm, 541 nm, 551 nm and 561 nm for CellTracker; 458 nm, 468 nm, 478 nm, 488 nm for actin-GFP). Following photobleaching, we recorded 200 post-bleach images every 0.23 s for both CellTracker and actin-GFP. For cytoplasm recovery measurements on *Plec*^+/+^ cells, we performed 38 measurements in 11 different cells, and 36 measurements on 11 cells for *Plec*^−/−^ cells. For actin turnover measurements, we performed 35 measurements in 15 different *Plec*^+/+^ cells and 27 measurements in 14 different *Plec*^−/−^ cells. Measurements were performed at varying locations distant from the nucleus and the cell membrane.

### Analysis of FRAP measurements

To determine the fluorescence recovery times for CellTracker and actin-GFP after photobleaching, we analyzed the time-dependent intensity during recovery *I* (*t*) in the bleached region throughout fluorescence recovery using a custom-written Python code. We computed the normalised intensity *I*_*n*_ (*t*) by dividing *I* (*t*) by the pre-bleach intensity *I*_*i*_. We corrected for bleaching by the probe beam by also measuring the fluorescence over time in a reference region far away from the bleaching region. Assuming a two-dimensional circular ROI and a uniform laser disk profile, we determined the FRAP recovery time by fitting *I*_*n*_ (*t*) to the following model (60):

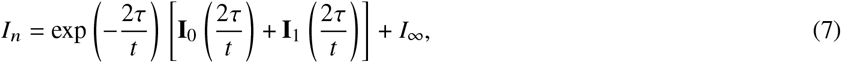

where *τ* is the characteristic recovery time, **I**_0_, **I**_1_ are modified Bessel functions and *I*_*∞*_ < 1 is the intensity asymptote due to an immobile fraction within the sample. For CellTracker, we can infer a diffusivity *D* from the relationship *τ* = *r*^2/^ (4*D*), with *r* being the bleach radius (60). It reacts with glutathione, which is most abundant in the cytoplasm, but is also present in subcellular organelles such as mitochondria (61). We thus expect to see partial rather than total fluorescence recovery, i.e., *I*_*∞*_ > 0. In the case of actin-GFP, we expect full fluorescence recovery (59).

### Western blotting

Cells were seeded in 6 well culture plates (Thermo Fisher Scientific, NC0506188) containing 2 mL of culture medium and allowed to attach for 48 hours. Cells were scraped in phosphate-buffered saline (PBS) and lysed in equal volumes of 2x Laemmli buffer (4% sodium dodecyl sulfate (SDS), 20% glycerol, 120 m M Tris(hydroxymethyl)aminomethane (Tris) buffer with pH 6.8). Lysates were cleared of large DNA by passing through a 25G needle and then heated to 65°C for 10 minutes. Protein concentrations were measured with the Lowry protein assay (62). Equal amounts of protein were size-separated by SDS-PAGE containing 2,2,2-Trichloroethanol (TCE). After running the gel, it was exposed to a 300 nm transilluminator (Bio-Rad ChemiDoc Imaging System) for 5 minutes. The separated proteins were then transferred to a polyvinylidene fluoride (PVDF) membrane (Bio-Rad, 1704156) using the Bio-Rad Trans-Blot Turbo Transfer System, and imaged on the transilluminator to get a picture of the total protein. Membranes were blocked for 1 hour at room temperature by 5% BSA in PBS supplemented with 0.1% Tween-20, incubated overnight at 4°C with primary antibody (see Supplementary Table 2). Subsequently, membranes were washed 5 times with 0.1% Tween-20 in PBS and then incubated with horseradish peroxidase-conjugated secondary antibodies (see Supplementary Table 2) for 1 hour at room temperature. After adding Enhanced chemiluminescence (ECL) substrate (Thermo Fisher Scientific, 32106), blots were imaged with the Bio-Rad ChemiDoc Imaging System. The band intensities were quantified using FIJI (57).

### Statistics

All mechanical experiments shown were repeated on at least 3 different days and imaging (including FRAP) experiments were repeated on 3 different days. Measured values in the main text are stated as the mean± the standard error on the mean. For all mechanical measurements, P-values to quantify statistical significance were calculated using the Wilcoxon rank-sum test after assessing normality with the Shapiro-Wilk test, which confirmed that the data did not follow a Gaussian distribution. For FRAP measurements, P-values were calculated using the unpaired Student’s t-test. Computer codes described in the Methods section are available on request.

## RESULTS

### Plectin influences actin and vimentin organization in fibroblasts

The aim of this study was to determine the contribution of plectin to the viscoelastic behaviour of fibroblasts at small and large deformations. To this end, we made a side-by-side comparison between immortalized mouse fibroblasts derived from transgenic plectin wild-type (*Plec*^+/+^) and plectin-deficient (*Plec*^−/−^) mice (for details see (43–45)). Western blot analysis confirmed that the *Plec*^−/−^ cells lack plectin, while the expression levels of actin and vimentin are comparable to wild type cells (Figure 1C and Figure S1). To measure the whole-cell mechanical response, we confined the cells between two parallel plates that were made nonadhesive to prevent any active response of the cells to compression.

**Figure 1:**
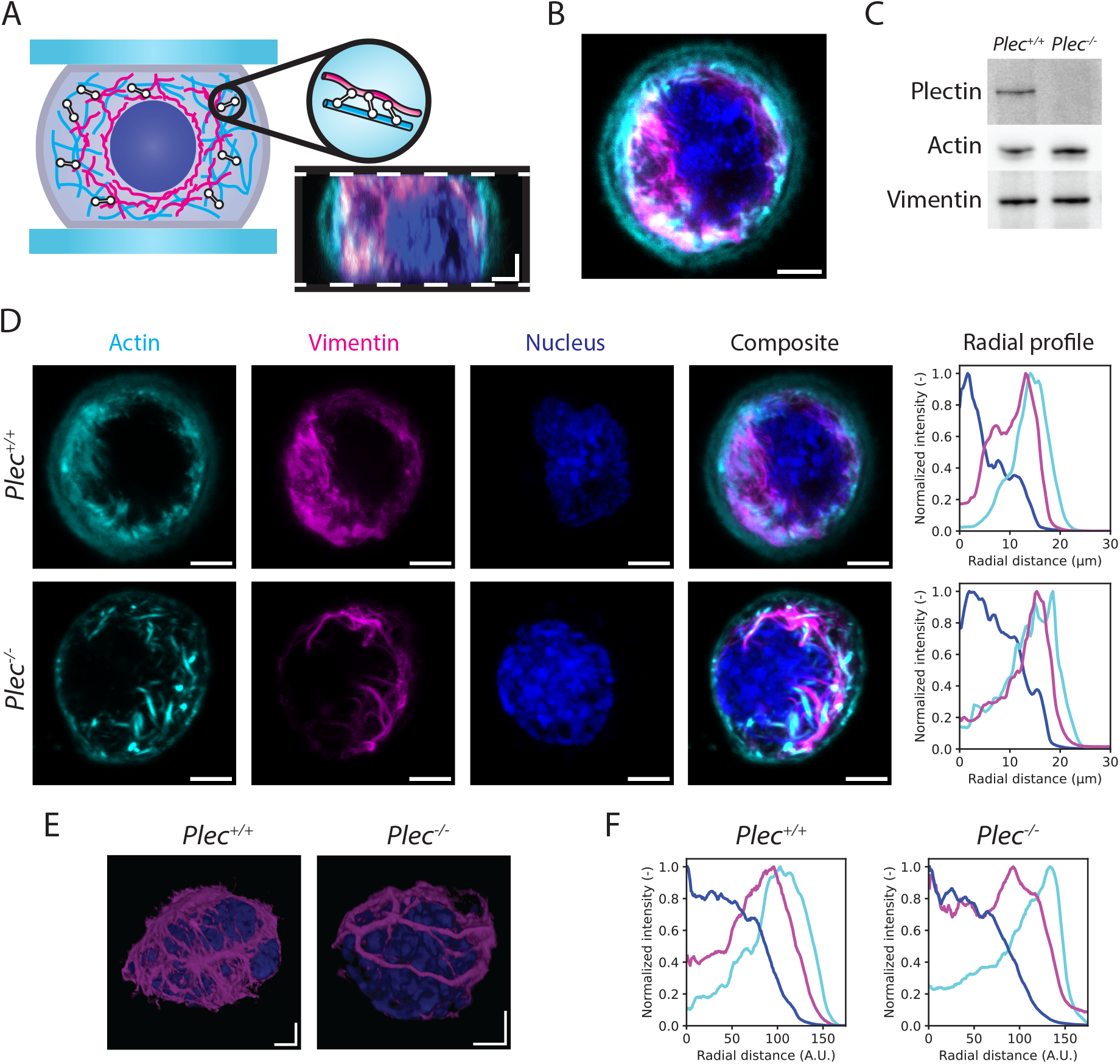
Plectin influences the organization of the actin and vimentin cytoskeleton in nonadherent fibroblasts. (A) The schematic shows how live fibroblasts were imaged while trapped between two nonadhesive surfaces. Plectin (white) forms crosslinks between actin filaments (cyan) and vimentin intermediate filaments (magenta). The image shows a side projection of a *Plec*^+/+^ fibroblast. White dashed lines indicate the two confining surfaces. Scale bars are 3 µm. (B) Single confocal slice through the cell equator of a *Plec*^+/+^ fibroblast. SiR-labeled actin (cyan) is concentrated at the cortex, while vimentin-GFP (magenta) surrounds the nucleus (dark blue). Scale bar 3 µm. (C) Immunoblot analysis of cell lysates from *Plec*^+/+^ and *Plec*^−/−^ cells using antibodies to plectin, actin and vimentin. (D) Representative images of *Plec*^+/+^ and *Plec*^−/−^ cells. Images are maximum intensity projections of 2 µm thick confocal sections at the cell equator. Actin is located largely at the cell periphery, while vimentin is located closer to the nucleus. Average radial profiles of the fluorescent signals (right) were calculated by taking normalized intensities around concentric circles from the center of the image. The radial distance is set to zero at the cell center. We observe a significant overlapping region between actin and vimentin. (E) 3D volume renderings of vimentin and the nucleus. In the *Plec*^+/+^ cell, a cage-like vimentin structure surrounds the nucleus, whereas in the *Plec*^−/−^ cell, vimentin forms bundles that no longer surround the nucleus on all sides. (F) We repeat the radial profile procedure shown in panel D by averaging the images of multiple cells (*N*=10 for both conditions) and computing the radial average. Average profiles resembled those of individual cells. Note that the scale of the ‘average cell’ images and line profile plots are in arbitrary units since the cells were rescaled to compensate for variations in cell size.

To characterize the impact of plectin knockout on the organization of the cytoskeleton in this geometry, we first imaged cells confined between two nonadhesive coverslips spaced apart by a distance of 10 µm (Figure 1A). This separation distance is smaller than the average cell size determined by imaging flow cytometry measurements (Figure S2), thus allowing us to image our cells in a compressed state. *Plec*^+/+^ and *Plec*^−/−^ cells had comparable diameters (22.4 ± 3.1 µm and 22.2± 3.1 µm, respectively, see Figure S2A) and nucleus sizes (8.3± 1.4 µm and 8.2 ± 1.4 µm, respectively, see Figure S2B). Side projections (Figure 1A) and top views (Figure 1B) reconstructed from confocal z-stacks showed that the cells are quasi-spherical with a radially rather uniform distribution of both actin and vimentin. Actin predominantly lines the plasma membrane at the cell periphery, but also extends into the cell interior (Figure 1D). By contrast, vimentin is primarily distributed between the actin-rich region and the nucleus (Figure 1D). Most strikingly, the vimentin network in the *Plec*^−/−^ cells displays increased bundling compared to the finer vimentin meshwork in *Plec*^+/+^ cells (second column in Figure 1D, compare top and bottom rows). This effect is strongly reminiscent of earlier observations of vimentin filament bundling in EBS-MD patient cells (28) and of keratin filament bundling in *Plec*^−/−^ keratinocytes (63). Furthermore, three-dimensional reconstructions of confocal z-stacks reveal that vimentin filaments in *Plec*^+/+^ cells are positioned in close proximity to the nuclear envelope, forming a filamentous network that encases the nucleus (Figure 1E, left). This structure has been referred to as the vimentin nuclear cage (64) and has been shown to protect the nucleus from mechanical stress (65). By contrast, the thicker vimentin bundles in *Plec*^−/−^ cells no longer form such a protective meshwork structure around the nucleus (Figure 1E, right).

To better compare the spatial distributions of actin and vimentin within the cell interior, we calculated radial distributions of the intensities. For individual cells, this showed a considerable overlap between the actin and vimentin localization (Figure 1D, right). To analyze this for many cells, we constructed ‘average cells’ by computing the average of maximum intensity projections from 2 µm thick confocal slices centered at the cell equator, adjusting for variations in cell size (Supplementary Figure 1). This analysis revealed similar nuclear distributions between *Plec*^+/+^ and *Plec*^−/−^ cells and confirmed a significant region of overlap between the actin and vimentin networks (Figure 1F). This observation suggests that the two networks interpenetrate. We observed subtle differences between *Plec*^+/+^ and *Plec*^−/−^ cells. The vimentin network on average extended to the cell periphery in *Plec*^−/−^ cells but not in *Plec*^+/+^ cells. The peak intensity for actin was close to the peak intensity for vimentin in *Plec*^+/+^ cells but closer to the cell periphery in *Plec*^−/−^ cells. We conclude that plectin has a significant impact on the organization of the actin and especially the vimentin cytoskeleton.

### Small amplitude oscillatory compression shows that plectin stiffens cells

To investigate how plectin removal impacts the mechanical response of the cells to small deformations, we used a single-cell compression device equipped with a wedged flexible cantilever. We subjected the cells to an oscillatory compressive strain with a small amplitude (∼ 1%) and stepwise increasing oscillation frequency from 0.1 Hz to 10 Hz (Figure 2A). From the resulting force on the cantilever, we calculated the real and imaginary parts of the frequency-dependent complex apparent

**Figure 2:**
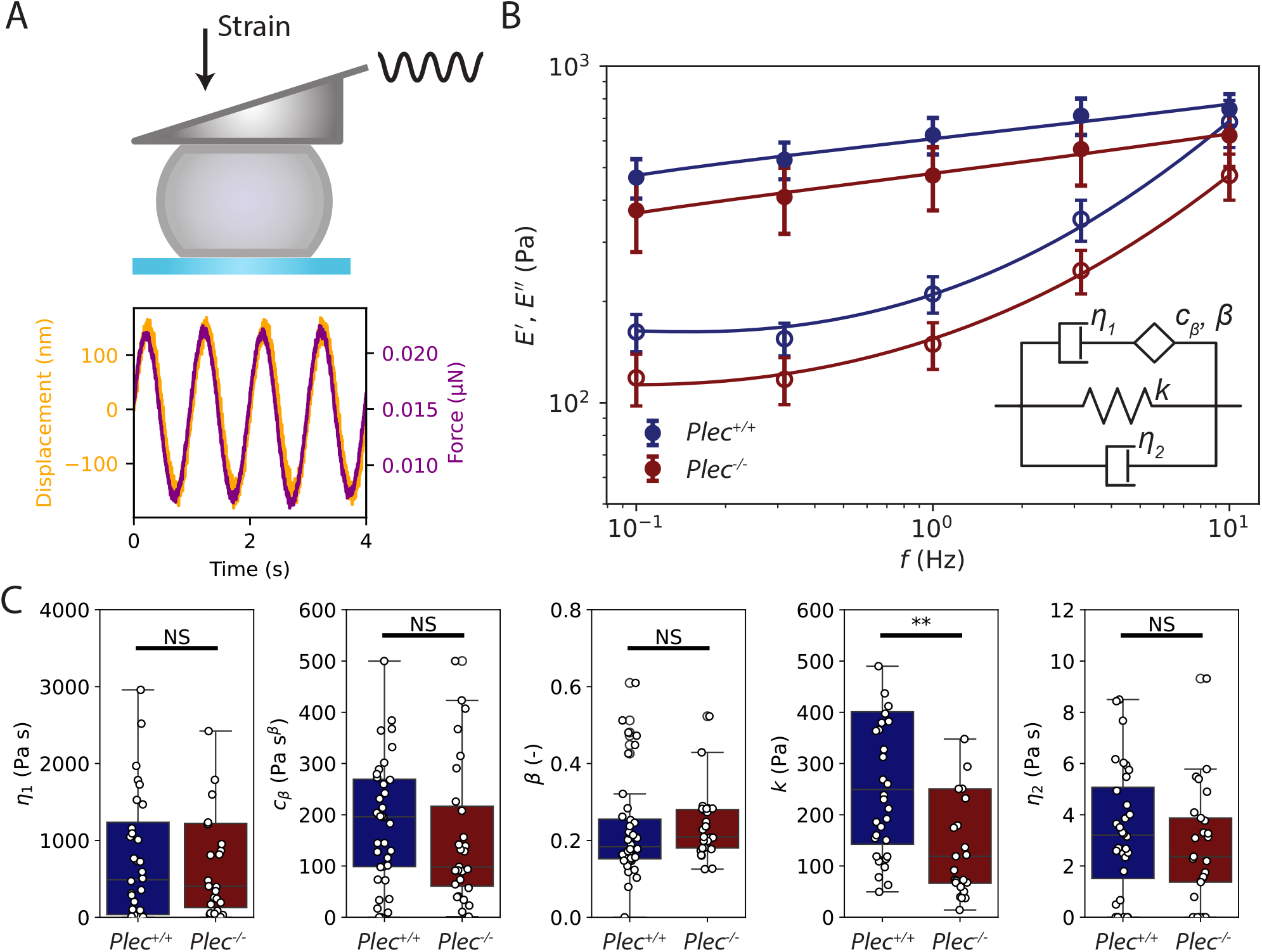
Small-strain oscillatory compression measurements on single fibroblasts reveal that *Plec*^+/+^ cells are stiffer than *Plec*^−/−^ cells. (A) Top: Illustration depicting a single cell positioned between a glass coverslip and the wedged cantilever. Bottom: Typical measurement for a *Plec*^+/+^ cell at an oscillation frequency of 1 Hz, showing the cantilever’s applied displacement (black; left y-axis) and the corresponding observed force (red; right y-axis) over time. (B) Storage (*E*^′^) and loss (*E*^′′^) linear compressive moduli (closed and open symbols, respectively), measured for oscillation frequencies between 0.1 Hz and 10 Hz for *Plec*^+/+^ cells (N=36, blue) and *Plec*^−/−^ cells (N=27, red). Fits on the average data (solid lines) to the fractional rheological model (inset) from Equation 2. (C) Material parameters determined from the fits of the fractional rheological model for each cell individually. The 5 fitting parameters are the dashpot viscosity *η* _1_ capturing long-term viscous dissipation, the spring-pot parameter *c*_*β*_ describing the intermediate viscoelastic behaviour, the fractional exponent *β* characterising power-law scaling with frequency, the elastic modulus *k* representing the stiffness of the material, and the Newtonian viscous term *η* _2_ capturing the response of the cytoplasm. Only the elastic parameter *k* changes upon plectin knockout, whereas all other parameters are unchanged. Asterisks indicates statistically significant differences (** denotes P<0.01), N.S. denotes non-significant differences.

Young’s modulus, *E*^*\**^ (*f*). The real part corresponds to the storage modulus *E*^′^ (*f*), which reflects the elastic component of the response (closed symbols in Figure 2B). The imaginary part corresponds to the loss modulus *E*” (*f*), which reflects the viscous component of the response (open symbols in Figure 2B). Both moduli show a qualitatively similar dependence on frequency for the *Plec*^+/+^ cells (blue) and *Plec*^−/−^ cells (red). In both cases, *E*^′^ is larger than *E*^′′^, in particular at low frequencies, consistent with predominantly solid-like behaviour. Furthermore, the elastic moduli increase with a weak power law dependence on frequency, characteristic for mammalian cells (66). At the highest frequency, the elastic and viscous moduli tend towards a crossover. In quantitative terms, however, we see a clear difference: The moduli of *Plec*^+/+^ cells are greater than those of *Plec*^−/−^ cells by a factor ∼1.4, indicating that plectin stiffens the cells. Additionally, we observe a shift in the projected crossover frequency for the elastic and viscous moduli, from ∼12 Hz in *Plec*^+/+^ cells to ∼16 Hz in *Plec*^−/−^ cells, signifying that *Plec*^−/−^ cells may maintain an elastic mechanical response over a broader frequency range.

For a more in-depth comparison of the *Plec*^+/+^ and *Plec*^−/−^ cells, we fitted the experimental data to phenomenological models that describe the rheology in terms of a small number of characteristic material parameters. We first attempted to describe the data in terms of the structural damping law, an established phenomenological model of cell rheology (50, 51).

This model (see Equation 1) has the benefit that it only has three material parameters, namely the scaling factor *E*_0_, power law exponent *α*, and cytoplasmic viscosity *μ*. We previously found that this model could capture the compressive modulus of mouse embryonic fibroblasts rather accurately (42). However, we found that it strongly underestimated the viscous modulus of the fibroblasts studied in this work, particularly at low frequencies (Figure S4). Apparently, we need to include an additional viscous contribution that captures slow stress relaxation. Therefore, we developed a new fractional rheological model inspired by recent work (52, 53) that combines short-term power-law stress relaxation and long-term exponential stress relaxation. This model (see Equation 2) has five free parameters: the elastic modulus *k* representing the stiffness of the material, the dashpot viscosity

*η*_1_ capturing long-term viscous dissipation, the spring-pot parameter *c*_*β*_ describing the intermediate viscoelastic behaviour, the fractional exponent *β* characterising power-law scaling with frequency, and finally the Newtonian viscous term *η*_2_ capturing the response of the cytoplasm. As shown in Figure 2B (solid lines), this model provides an accurate and self-consistent fit of *E*^′^ and *E*^′′^. The only fitting parameter that is significantly different between the *Plec*^+/+^ and *Plec*^−/−^ cells is the elastic modulus *k*, which decreases from *k* = 232± 20 Pa in *Plec*^+/+^ cells (N=36) to *k* = 120 ±20 Pa in *Plec*^−/−^ cells (N=27) (Figure 2C). The two parameters characterizing the power-law scaling behavior, *β* and *c*_*β*_, are comparable between *Plec*^+/+^ and *Plec*^−/−^ cells. Likewise, the dashpot viscosity *η*_1_ and Newtonian damping term, *η*_2_, were unchanged. We conclude that plectin only affects the elastic properties of the cytoskeleton at small deformations, within the frequency range probed here.

### Plectin affects poroelastic and viscoelastic stress relaxation

The oscillatory compression measurements suggest that there is significant energy dissipation on time scales beyond 10 seconds, i.e., at frequencies below the measurement limit of 0.1 Hz. Since many organs and connective tissues experience deformations that persist for minutes (67, 68), we decided to measure the viscoelastic behaviour of the *Plec*^+/+^ and *Plec*^−/−^ cells on longer timescales via step strain experiments (Figure 3A top). We subjected cells to a rapid compression by a distance of 1 µm, corresponding to a small (∼ 0.05) strain within the linear elastic regime. We then measured the stress relaxation dynamics whilst holding this displacement for 80 s (Figure 3A bottom). As shown in Figure 3B, both *Plec*^+/+^ cells (blue solid line) and *Plec*^−/−^ cells (red solid line) display marked stress relaxation over this time scale (for individual cell data see Figure S5). The stress is overall larger for the *Plec*^+/+^ cells as compared to the *Plec*^−/−^ cells, indicating that plectin depletion causes cell softening, consistent with the small amplitude oscillatory data. In a log-log representation, we can distinguish multiple relaxation regimes. At short times (< 0.1 s, indicated by a grey background in Figure 3B), we observe an exponential stress decay. Here, the mechanical response to compression is known to be dominated by the flow of cytosol through the porous elastic structure provided by organelles and the cytoskeleton (59) (see schematic inset of Figure 3B). Beyond 0.1 s, the stress relaxation response transitions to a power-law time dependence, reflecting a broad distribution of relaxation processes. In this regime we indeed expect power-law viscoelastic stress relaxation just like in the small amplitude oscillatory measurements. This is followed by exponential relaxation at long times.

**Figure 3:**
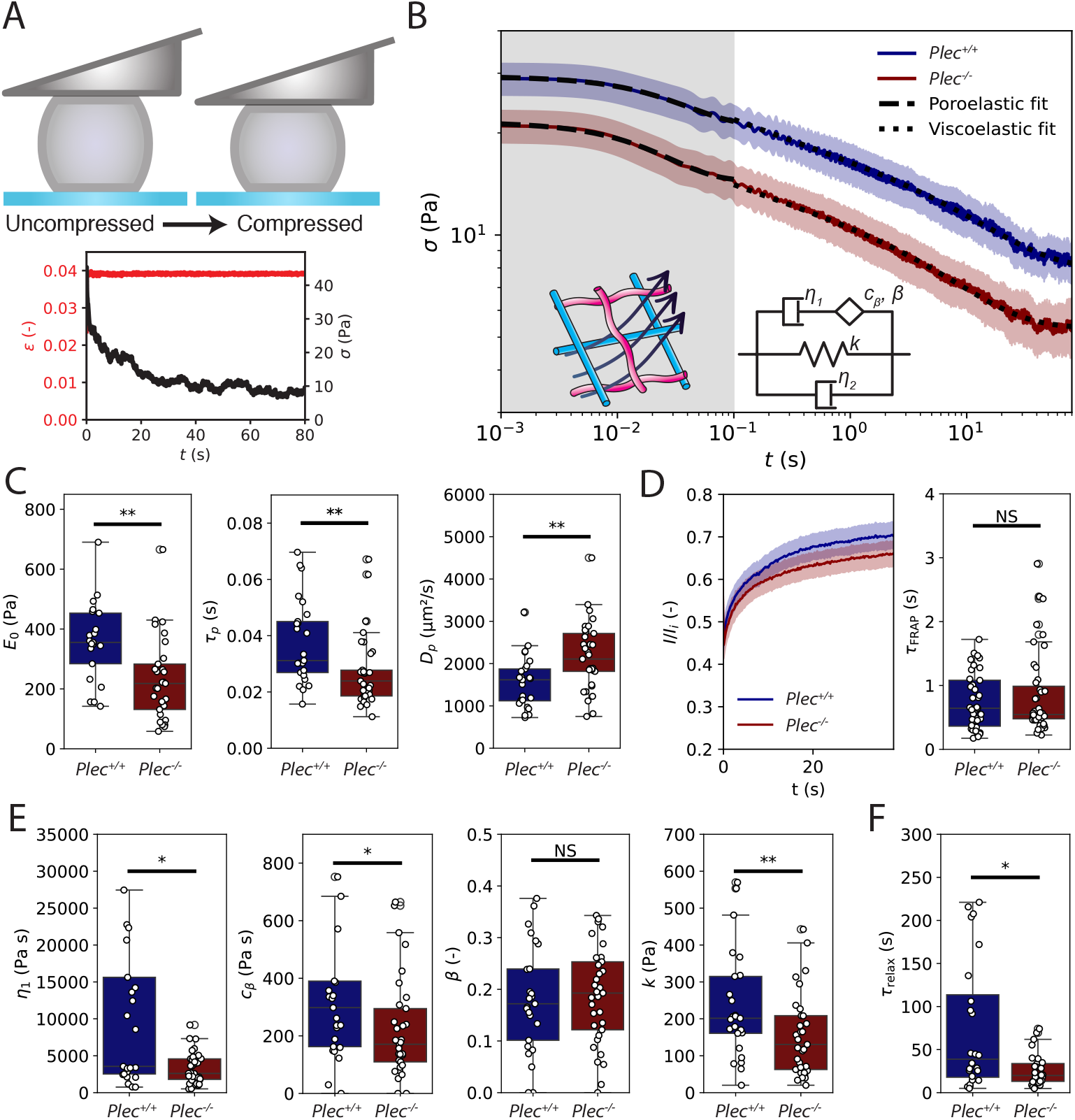
Small amplitude stress relaxation experiments for *Plec*^+/+^ and *Plec*^−/−^ cells. (A) Top: Schematic of the step-strain experiment (not to scale, the strain is only 0.05). Bottom: Typical measurement for a *Plec*^+/+^ cell showing the cantilever displacement that is held constant (red; left y-axis), while the force (black; right y-axis) is measured over time. (B) Averaged stresses for *Plec*^+/+^ cells (blue solid line, N = 25) and *Plec*^−/−^ cells (red solid line, N = 33) over time in log-log representation. The shaded areas around the average relaxation curves represent the standard deviation. Stresses are higher for *Plec*^+/+^ cells, indicating these cells are stiffer. At short time scales (from 0.001 s to 1 s, indicated by the grey background), the cells display exponential stress relaxation characteristic of a poroelastic material, where fluid flows through the cytoskeletal network (inset drawing). Fits to the poroelastic model (equation 5) are shown as dashed lines. At later times (from 0.1 s to 80 s, white background) the data are well-described by the same fractional rheological model (inset schematic) that also captures the oscillatory data, transformed to the time domain (fits shown as dotted lines; equation 6). (C) Elastic moduli *E* _0_ (left) and poroelastic time scales *τ*_p_ (middle) determined from the poroelastic model fits for individual cells. The poroelastic diffusion coefficient *D*_p_ (right) calculated from *τ*_*p*_ data is faster for *Pl ec*^−/−^ cells than for *Plec*^+/+^ cells. (D) Comparison of cytoplasmic diffusion via fluorescence recovery after photobleaching (FRAP). Left: average FRAP recovery curves for *Plec*^+/+^ and *Plec*^−/−^ cells. The error band shows the standard error. Right: We measure no significant difference between the FRAP recovery times, *τ*_FRAP_, of *Plec*^+/+^ and *Plec*^−/−^ cells. (E) Material parameters obtained from the fractional rheology fits, showing the dashpot viscosity *η*_1_ describing long-term viscous dissipation, the spring-pot parameter *c*_*β*_ describing the intermediate viscoelastic behaviour, the fractional exponent *β* characterising power-law scaling with frequency, and the elastic modulus *k* representing cell stiffness. (F) Viscoelastic stress relaxation time scale *τ*_relax_ calculated from the data in panel E using Equation 10. Asterisks indicate statistically significant differences (* denotes P<0.05, ** P<0.01), N.S. denotes non-significant differences.

To test how plectin changes the poroelastic and viscoelastic response of the cells, we fitted the stress relaxation data in the short and intermediate time regimes to corresponding theoretical model predictions. The short-time poroelastic response is expected to be single-exponential (Equation 5) with a characteristic poroelastic decay time *τ*_p_, assuming that the cell can be approximated as a simple biphasic material consisting of a porous elastic meshwork with an elastic modulus, *E*_0_, bathed in an interstitial fluid (cytosol) of viscosity *μ*. The data were well-described by this model (dashed lines in Figure 3B). From the fits, we found that plectin knockout significantly softens the cells, with an instantaneous elastic modulus (stiffness of the cell at t=0) of *E*_0_ = 230± 30 Pa for *Plec*^−/−^ cells compared to *E*_0_ = 360 ±30 Pa for *Plec*^+/+^ cells (Figure 3C left). Furthermore, poroelastic relaxation occurs more rapidly upon plectin knockout, with a poroelastic time scale of *τ*_p_ = 0.040± 0.001 s in *Plec*^−/−^ cells as compared to *τ*_p_ = 0.030± 0.001 s in *Plec*^+/+^ cells (Figure 3C middle).

The difference in poroelastic time scales suggests that the porositity of the cytoplasm might be higher in *Plec*^−/−^ cells compared to *Plec*^+/+^ cells. To test this idea, we first estimated the poroelastic diffusion constant *D*_p_ from the poroelastic time scales according to:

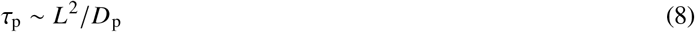

For simplicity, we assume that the elastic network of cytoskeleton and organelles is homogeneous across the distance between the cell membrane and nuclear envelope so we can define the characteristic length scale *L* as the distance between the cell membrane and the nuclear envelope. We can estimate this distance from the earlier mentioned flow cytometry measurements of the cell and nucleus sizes (Figure S2). *Plec*^+/+^ and *Plec*^−/−^ cells had varying sizes but both displayed a proportional increase of cell size with nucleus size (Figure S2E,F), resulting in a rather constant nucleus-to-cell diameter ratio of ∼0.74 in both cases. We therefore assume a characteristic length scale of *L* = 7.1µm and thus estimate *D*_p_ = (1500 ±100) µm /s for *Plec*^+/+^ cells and *D*_p_ = (2200 ±100) µm^2^/s for *Plec*^−/−^ cells (Figure 3C right). Note that if we assume fluid transport also occurs in the nucleus, so *L* equals the cell size, we still find a larger *D*_p_ in *Plec*^−/−^ cells (see supplementary information for more details). The poroelastic diffusion constant *D*_p_ depends on the pore radius *ξ* of the cytoskeletal network, the cytosol viscosity *μ*, and the elastic modulus of the drained cytoskeletal network *E* according to (59):

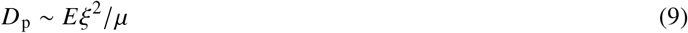

To measure the unknown cytosol viscosity and test whether it changes upon plectin knockout, we performed fluorescence recovery after photobleaching (FRAP) on the cytoplasmic dye CellTracker Orange. We photobleached a small (1 µm) circular region and observed the recovery of the fluorescent signal (Figure 3D and Figure S6). By fitting the individual recovery curves to Equation 7, we recovered comparable recovery times for *Plec*^+/+^ cells (*τ*_FRAP_ = 0.53 ± 0.05s) and *Plec*^−/−^ cells (0.50 ± 0.06s), indicating comparable cytosol viscosities. We estimate a cytoplasmic viscosity *μ* ≃ 4 Pa s for both *Plec*^+/+^ and *Plec*^−/−^ cells using the Stokes-Einstein relation for the diffusion of small molecules 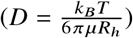 (69), assuming a hydrodynamic radius *R*_*h*_ ≃ 0.5 nm (70) for the cytoplasmic dye. Encouragingly, this viscosity value is consistent with the cytoplasmic viscosity (*η*_2_ = 3.3± 0.4) Pa s) inferred by fitting the oscillatory compression data to the fractional rheological model (Figure 2C), lending support for this model. Plugging the poroelastic diffusion times from the step-strain data and the cytosol viscosity from the FRAP data into Equation 9, we calculate that the cytoskeletal meshwork of *Plec*^−/−^ cells (*ξ* = 7.2µm) has larger pores than for *Plec*^+/+^ cells (*ξ* = 4.9µm) by a factor of ∼ 1.5. Interestingly, this is consistent with the observation that the plectin knockout cells have a more open, bundled vimentin cytoskeleton than wild type cells (Figure 1D) while having similar concentrations of vimentin and actin (Figure 1C).

We next turned to the viscoelastic stress relaxation regime at intermediate times (0.1 s to 80 s), which should be describable by the same fractional rheological model used to model the oscillatory measurements (inset showing a schematic of the model in Figure 3B). To test this, we converted the model from the frequency domain (Equation 2) to the time domain (Equation 6), enabling a direct comparison of the characteristic mechanical parameters. Note that the model involves 4 fit parameters in the time domain instead of 5 parameters in the frequency domain, since the 5th parameter, *η*_2_, is reduced to a Dirac delta function after the Laplace transform. The fractional rheology model indeed fitted the force–relaxation curves well (dotted black lines in Figure 3B), returning similar conclusions as the oscillatory measurements. Again, we observed a significant decrease in the elastic modulus *k*, from *k* = 250 ±30 Pa for *Plec*^+/+^ cells to *k* = 170± 30 Pa in *Plec*^−/−^ cells (Figure 3E) but an unchanged power-law scaling exponent *β*. However, contrary to the oscillatory measurements, the viscous modulus *η*_1_ determined from the stress relaxation significantly decreased upon plectin knockout, from *η*_1_ = 8000 ±2000 Pa for *Plec*^+/+^ cells to *η*_1_ = 3700± 500 Pa for *Plec*^−/−^ cells. The stress relaxation data are likely more accurate here because we could measure up to time scales of 80 s, much longer than in oscillatory measurements. We can calculate the viscoelastic relaxation time, *τ*_relax_, that marks the transition from a power-law decay at intermediate times to an exponential relaxation regime at longer times inaccessible in the oscillatory measurements according to (53):

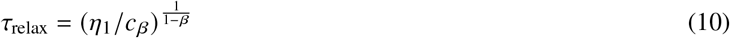

This timescale indicates when the dashpot characterizing long-time scale relaxation processes (beyond ∼ 10s) in the cytoskeleton begins to dominate stress relaxation (71). We observed a statistically significant decrease in this time scale, from *τ*_relax_ = 75 ±15 s for *Plec*^+/+^ cells to *τ*_relax_ = 26± 4 s in *Plec*^−/−^ cells, indicating faster stress relaxation upon plectin knockout (Figure 3F).

On long time scales, we expect that stress relaxation could occur due to unbinding of plectin and other cytoskeletal crosslinker proteins or by cytoskeletal turnover. Actin filaments turn over as a consequence of filament depolymerization, which is catalyzed by various actin-binding proteins. Actin network turnover times are typically reported to be on the order of seconds to minutes (72, 73), which could correspond to the time scale where we observe stress relaxation. The vimentin cytoskeleton is typically reported to turnover on much longer time scales on the order of hours, via subunit exchange along the filaments and fragmentation (74). To determine the actin and vimentin turnover rates in our cells and test whether they are changed by plectin knockout, we performed fluorescence recovery after photobleaching (FRAP) experiments on cells tagged with GFP-actin (Figure 4A) and GFP-vimentin (Figure S8). As shown in Figure 4B, the GFP-actin signal averaged over many cells fully recovered within 60 s for both *Plec*^+/+^ and *Plec*^−/−^ cells (blue and red curves, respectively; see Figure S7 for individual curves). Strikingly, however, the actin turnover rate as quantified by the FRAP recovery time by fits to equation 7 was faster in *Plec*^−/−^ cells by a factor of ∼ 3 (Figure 4C). By contrast, FRAP experiments on GFP-vimentin transfected cells showed negligible recovery up to times of 80 s (corresponding to the time scale probed in the step strain experiments) for both *Plec*^+/+^ and *Plec*^−/−^ cells (Figure S8). We therefore conclude that the faster viscoelastic stress relaxation rate in the plectin knockout cells at long times is likely due to faster actin turnover.

**Figure 4:**
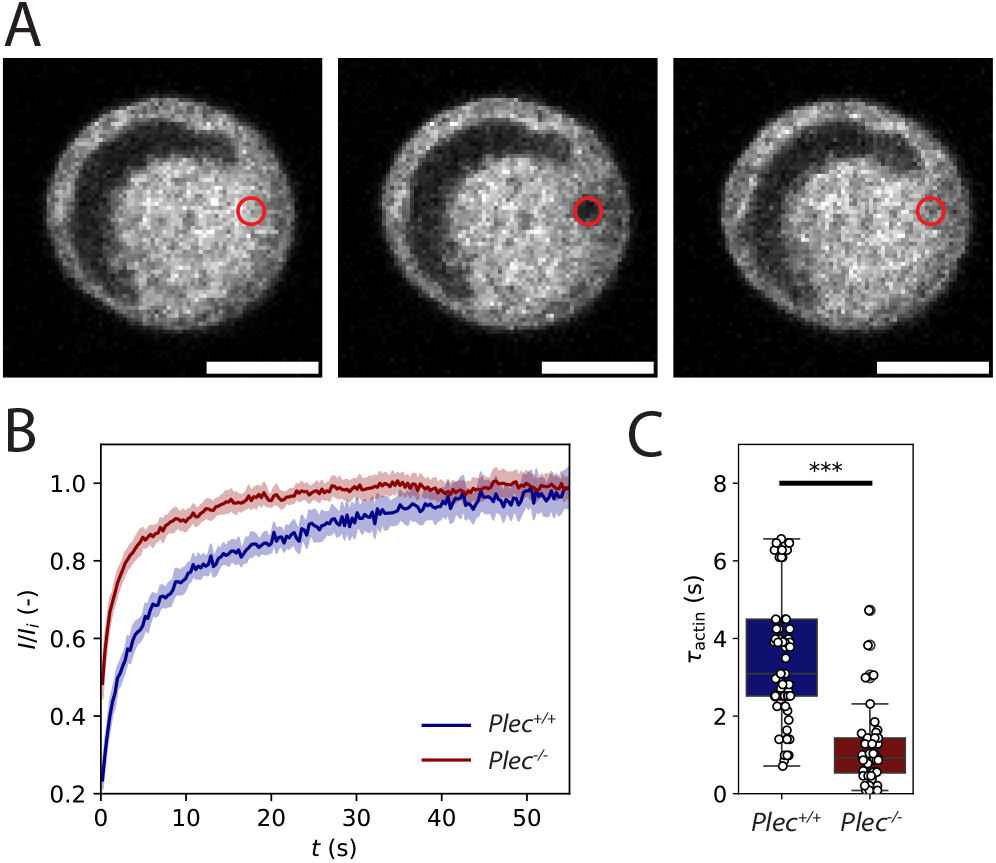
Comparison of actin turnover rates in *Plec*^+/+^ and *Plec*^−/−^ cells confined between two nonadhesive surfaces via fluorescence recovery after photobleaching (FRAP) experiments. (A) We bleach a small (1 µm) circular region (red open circle) in a region of the cell where we observe an actin network (tagged with GFP-actin) and monitor the fluorescence recovery over time. Note that the large dark region within the cell is occupied by the nucleus. Scale bar 10 µm. (B) Averaged FRAP recovery curves for GFP-actin-tagged *Plec*^+/+^ cells (blue, N = 54) and *Plec*^−/−^ cells (red, N = 35). In both conditions, the fluorescence fully recovers, but with a significantly higher rate in the *Plec*^−/−^ cells. (C) FRAP recovery times *τ*_actin_ obtained by fitting to the individual recovery curves (shown in Supplementary Figure S7). We find average values of *τ*_actin_ = 3.6± 0.4s for *Plec*^+/+^ cells and 1.4± 0.2 for *Plec*^−/−^ cells, indicating ∼ 3-fold faster actin turnover upon plectin knockout. *** indicates statistically significant differences (P<0.001).

### Plectin knockout does not impact fibroblast strain-stiffening

To test the effect of plectin on the mechanical response of fibroblasts to large compressive deformations, we performed strain ramp experiments (Figure 5A). The cells were compressed gradually with a constant rate of 1 µm/s, which is slow enough to minimise stress from viscoelastic contributions (42). Upon reaching the target deformation (*ϵ* ∼ 0.4), the cantilever was immediately retracted at the same rate. Figure 5B shows an example stress-strain cycle performed on a *Plec*^+/+^ cell. The stress-strain curves during compression (indicated by the arrow up) and retraction (arrow down) do not overlap: for any given strain, the corresponding stress is lower during retraction than it is during compression. This hysteresis confirms the viscoelastic nature of the cells. We next compared strain ramp experiments on *Plec*^+/+^ cells (N = 13) and *Plec*^−/−^ cells (N = 15) (see Figure S9 for individual stress-strain curves). The averaged data for the response upon compression reveal that *Plec*^−/−^ cells are softer than *Plec*^+/+^ cells: at any strain, the average resulting stress is lower for *Plec*^−/−^ cells (red) than for *Plec*^+/+^ cells (blue) (Figure 5C). However, the stress in both cases increases non-linearly with the applied strain. This type of compression-stiffening behavior under uniaxial load is consistent with earlier studies (42, 75).

**Figure 5:**
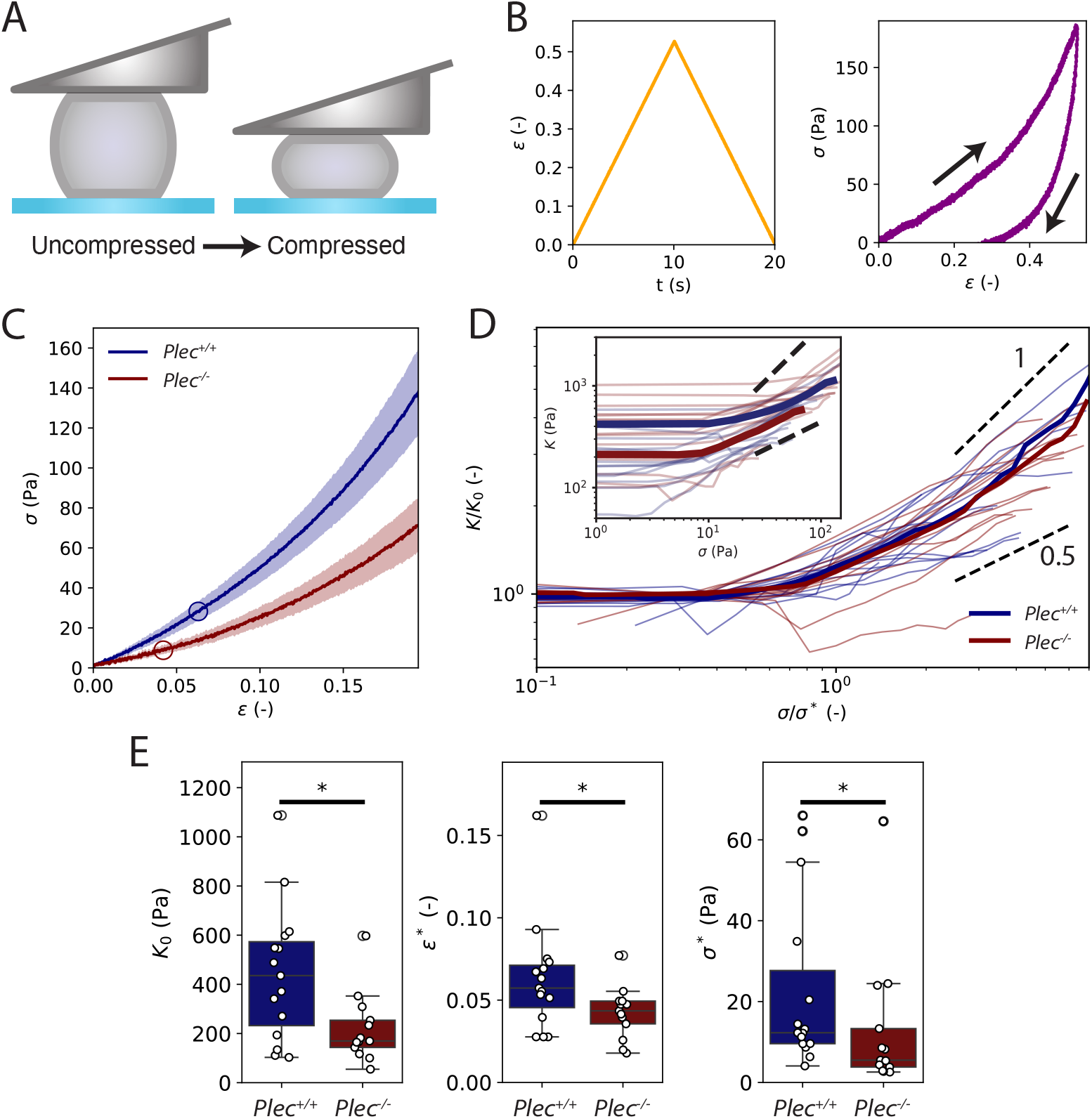
Strain ramp measurements of the nonlinear elastic response of *Plec*^+/+^ and *Plec*^−/−^ cells to large uniaxial compressions. (A) Schematic of the experiment where cells are compressed by gradually lowering the cantilever at a constant rate until a target strain of ∼ 0.4, and then unloaded by raising the cantilever back to its original zero-strain position. (B) Representative strain ramp measurement of a single *Plec*^+/+^ cell, showing the applied strain (left) and concomitant stress (right). The arrows show the order of loading and unloading. (C) Average stress-strain curves for *Plec*^+/+^ (N=15) and *Plec*^−/−^ (N=13) cells showing that *Plec*^−/−^ cells are softer than *Plec*^+/+^ cells but both cell types stress-stiffen. Shaded areas around the curves show the standard error. Circles indicate the onset of strain-stiffening. (D) Differential elastic modulus, *K*, for individual cells, obtained by taking the numerical derivative of the stress-strain curves from individual cells shown in Supplementary Figure 9. Inset shows the raw data, main plot shows the same data with *σ* normalized by the onset stress *σ*^*\**^ and *K* normalized by the linear modulus *K*_0_. *Plec*^−/−^ cells attain a similar stiffness as *Plec*^+/+^ cells at large stress (inset) and normalization collapses the curves onto a single master curve. The short dashed lines indicate that the stress dependence falls between power-law dependences with an exponent between 0.5 and 1. (E) Comparison of the stress-stiffening behavior of *Plec*^+/+^ cells and *P lec*^−/−^ cells in terms of the linear modulus, *K*_0_ (left), onset strain, *ϵ* ^*\**^ (middle), and onset stress, *σ*^*\**^ (right). *Plec*^−/−^ cells (red) are about two-fold softer than *Plec*^+/+^ cells (blue) and, accordingly, display a smaller onset stress. * indicates statistically significant differences (P > 0.05).

To quantify this strain-stiffening behaviour, we defined the differential stiffness *K* as *K* = d*σ*/ d*ϵ*. The stiffness is constant (*K* = *K*_0_) up to a certain onset strain *ϵ* ^*\**^, above which stiffening occurs (Figure 5D inset). Strikingly, the stiffness at large compressive stresses of 80 Pa is comparable for *Plec*^+/+^ cells (blue) and *Plec*^−/−^ cells (red). Also, normalizing the stiffness by *K* = *K*_0_ and the stress by *σ*^*\**^ collapses the data onto a single master curve (Figure 5D). At large deformations, the stiffness approaches a power-law dependence on stress with best-fit values for the power law exponent being similar for *Plec*^+/+^ cells (0.49± 0.06) and *Plec*^−/−^ cells (0.49 ±0.05). We note that the nonlinear regime covers less than a decade in stress, so these values should be taken as indicative only. The short dashed lines in Figure 5D show that the exponents lie somewhere between and 1.

As shown in Figure 5E (left), the linear modulus *K*_0_ is about 2-fold higher for the *Plec*^+/+^ cells (440± 70 Pa) as compared to the *Plec*^−/−^ cells (220± 40 Pa). This effect is consistent with the 1.7-fold difference found by small amplitude oscillatory and step strain measurements. The onset stress for stress-stiffening (*σ*^*\**^) is also higher for *Plec*^+/+^ cells (22± 5Pa) as compared to *Plec*^−/−^ cells (13 ±5Pa), consistent with their higher modulus (Figure 5E, right). The corresponding onset strains for strain-stiffening (*ϵ* ^*\**^) are therefore only slightly different between *Plec*^+/+^ cells (0.063 ±0.008) versus *Plec*^−/−^ cells (0.042 ± 0.004) (Figure 5E, middle).

### Repeated compression softens wild type cells more than plectin knockout cells

In many organs, such as the heart, skin and lungs, cells are subject to repeated compression. To test whether plectin knockout changes the mechanical response of fibroblasts upon repeated loading, we applied five subsequent linear compression-decompression strain ramps with an amplitude of 5 µm and strain rate of 1 µm/s (Figure 6A). Figure 6B shows a typical example of the strain (yellow) and stress (purple) for a *Plec*^+/+^ cell. In each subsequent cycle, we see that the stresses decrease for every given strain. To rule out the possibility that this softening was due to cell flattening, we performed widefield fluorescence imaging to monitor for possible changes in cell diameter or other morphological features. We found that the cell diameter returned to its original value after each cycle, indicating that no gross shape changes occurred with repeated compression (Figure S10). The cell contours remained smooth as long as the strain remained below 0.5, beyond which point cells often started to bleb as a sign of damage.

**Figure 6:**
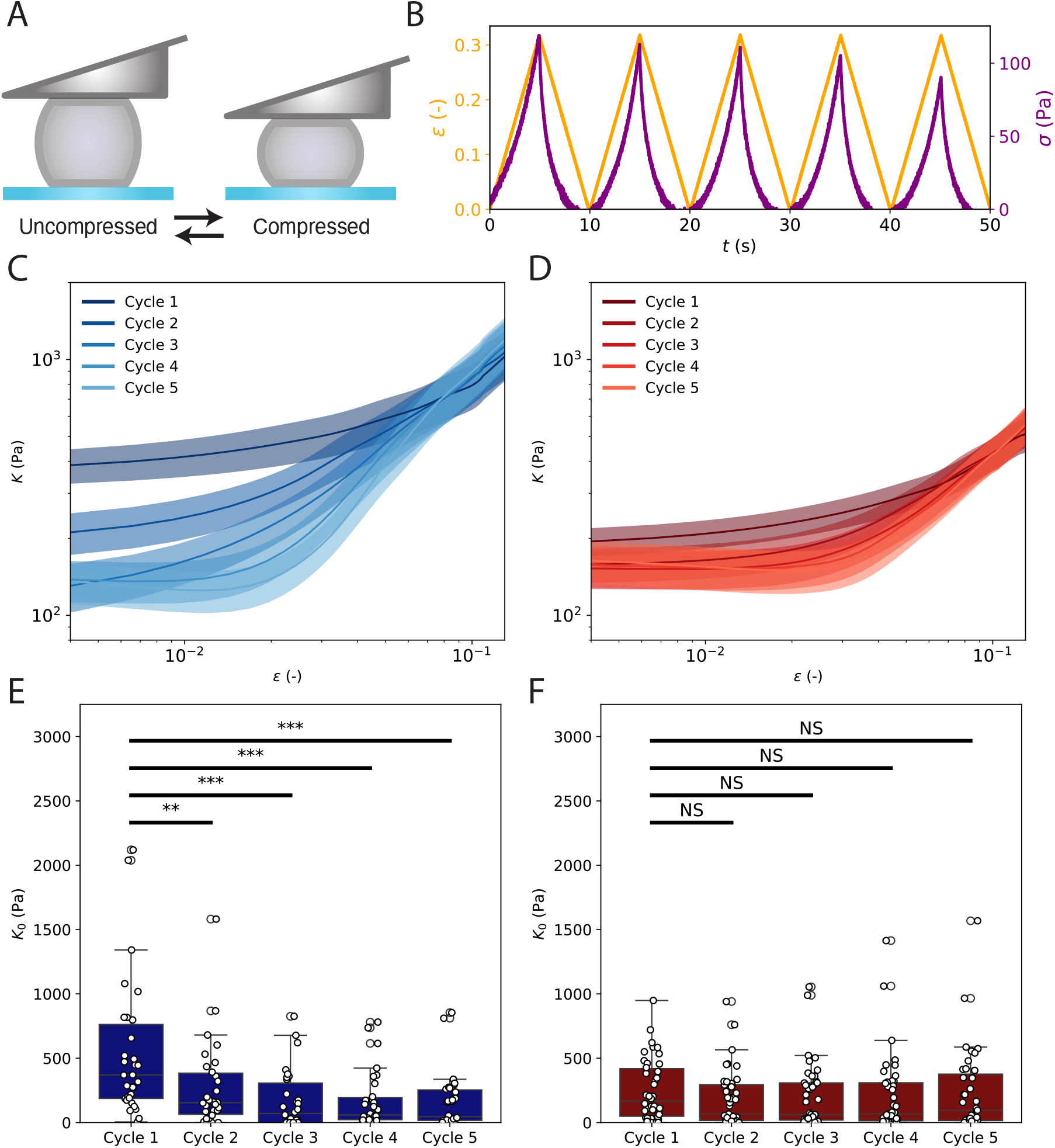
Mechanical response of *Plec*^+/+^ and *Plec*^−/−^ cells to repeated compressive loading-unloading cycles. (A) Schematic showing toggling between an uncompressed state (zero strain) and compressed state (strain ∼ 0.4) by gradual lowering and raising of the cantilever. (B) Example measurement on a *Plec*^+/+^ cell showing the applied strain (yellow, left y-axis) and measured stress (purple, right y-axis). For each subsequent cycle, the stress decreases. (C) Averaged dependence of the differential modulus *K* on applied strain for *Plec*^+/+^ cells (N = 30). We observe a large reduction in *K* at small strain between the first and fifth cycle of compression, whereas the curves converge at high strain. (D) Average *K*/ *ϵ* curves for *Pl ec*^−/−^ cells (N = 42), showing little change in the modulus between cycles. Error bands in panels C and D represent the standard error. (E) Linear modulus *K*_0_ as a function of cycle number for *Plec*^+/+^ cells obtained from fitting data for individual cells (shown in Supplementary Figure 9). (F) *K*_0_ as a function of cycle number for *Plec*^−/−^ cells. Note that the *Plec*^+/+^ cells show significant softening upon repeated loading (** denotes P<0.01, *** denotes P<0.001), while the *Plec*^−/−^ cells do not soften appreciably (differences N.S.).

To quantify the degree of softening upon repeated loading-unloading cycles, we calculated the differential stiffness *K* for each cycle of deformation, following the same procedure as for the single cycle compressive ramps. For *Plec*^+/+^ cells we found a substantial reduction in cell stiffness in the linear regime between the first and second cycle, with a further gradual reduction as cells were subjected to additional cycles (Figure 6C). At high stress, the stress-strain curves converged. For *Plec*^−/−^ cells we found qualitatively similar behavior, but the reduction of the stiffness in the linear regime between the first and subsequent cycles was comparatively minor (Figure 6D). To better visualize this, we tracked the value of the linear modulus *K*_0_ as a function of cycle number. For *Plec*^+/+^ cells, *K*_0_ was 400± 70 Pa in the first cycle and only 120± 30 Pa in the fifth cycle (Figure 6E). By contrast, *Plec*^−/−^ cells displayed a comparable linear modulus throughout the cycles, with average *K*_0_ values of 180 ± 30 Pa for the first cycle and 160± 40 Pa for the fifth cycle (Figure 6F). Interestingly, these stiffness values are comparable to the stiffness of *Plec*^+/+^ cells after repeated compression, strongly suggesting that the plectin crosslinkers that contribute to the modulus initially are disrupted by large compressive deformations.

## DISCUSSION

The aim of this study was to determine the contribution of plectin to the viscoelastic response of fibroblasts at small and large deformations using a custom single-cell rheometer. We first assessed the linear viscoelastic response using two complementary assays: small-amplitude oscillatory compressions and stress relaxation measurements. Both assays independently confirmed that fibroblasts behave as viscoelastic solids, displaying an elastic Young’s modulus, *E*^′^ (*f*), that is larger than the loss modulus, *E*” (*f*), with weak dependencies of the moduli on frequency. We next applied large uniaxial strains to investigate the nonlinear elasticity of the cells. Under these conditions, the cells showed pronounced strain-stiffening, with the differential elastic modulus rising sharply beyond ∼ 4% compressive strain. Both the small and large amplitude experiments showed that plectin knockout makes the mouse embryonic fibroblasts about 2-fold stiffer. This finding is in line with previous measurements on mouse skin fibroblasts (35) and mouse myoblasts (37). These studies used localized measurement methods in contrast to our whole-cell compression experiments, but also found *Plec*^−/−^ cells to be softer than *Plec*^+/+^ cells (see Table 1). However, other studies reported a modest 10% increase in stiffness upon plectin knockout in mouse keratinocytes (37) and no significant change in vulvar carcinoma cells (38). These contradictory findings could stem from variations in cell lines and cell types used and/or from the different measurement techniques employed.

**Table 1:**
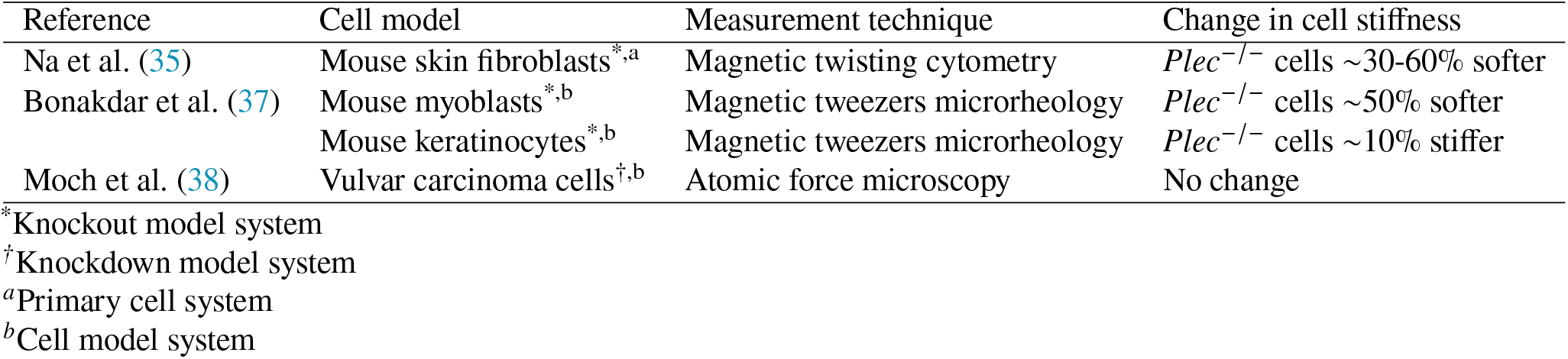
Summary of previously reported changes in the stiffness of single cells at small deformations upon plectin knockout for different cell model systems and with different measurement techniques.

A more detailed interpretation of our data aided by available theoretical models showed intriguing effects of plectin on the time- and amplitude-dependent mechanical response of fibroblasts. First, the stress relaxation measurements revealed that plectin reduced the poroelastic time scale that characterizes stress relaxation via intracellular fluid flow at short times (< 1 s) after a rapid compression (59). By also measuring the cytosolic viscosity by FRAP measurements on a small cytoplasmic dye, we found that this effect is caused by a larger pore size of the solid phase of the cytoplasm. The solid phase comprises the cytoskeleton, nucleus, and other organelles. Confocal imaging and flow cytometry showed that the wild type and plectin knockout cells had similar sizes and also similarly sized nuclei. However, the organization of the vimentin network was drastically different, with a dense meshwork in wild type cells versus a sparse network of bundles in plectin knockout cells. This bundling is expected to increase the pore size of the cytoskeletal matrix, which should indeed enhance the rate of poroelastic relaxation. Our data therefore suggest that plectin impacts poroelastic relaxation by regulating the organization of the vimentin cytoskeleton. This could be a consequence of its crosslinking role, both as a vimentin-vimentin and vimentin-actin crosslinker. In addition, anchoring of the vimentin network to focal adhesions and the nucleus by plectin could also help prevent vimentin filaments from collapsing into bundles.

A second finding from the stress relaxation experiments was that viscoelastic stress relaxation at times beyond the poroelastic regime obeys a fractional rheology model. According to this model, the cells display power law rheology for frequencies between 0.1 Hz and 10 Hz, characterized by a weak power law dependence of *E*^′^ (*f*) on *f*. Power law rheology signifies a broad distribution of characteristic time scales and is typical of mammalian cells (50). On longer time scales, the model includes an additional stress relaxation mechanism characterized by an exponential dependence on time. The same fractional rheology model could also self-consistently capture the frequency dependence of both the elastic and viscous moduli measured through oscillatory measurements. This consistency suggests that this model, though phenomenological, provides an adequate description of the main mechanisms contributing to the cell’s viscoelastic response. By fitting the data, we could compare the corresponding material parameters between *Plec*^+/+^ and *Plec*^−/−^ cells.

We found that removal of plectin altered the elastic component of the complex modulus, marked by a reduction in the elastic parameter *k*. It did not significantly change the power law scaling exponent *β* describing the frequency dependence at intermediate times. This result indicates that plectin’s crosslinking function predominantly reinforces the elastic integrity of the cytoskeletal network, while the fundamental mechanisms governing energy dissipation at intermediate timescales remain intact. This is consistent with previous studies on reconstituted actin networks with cross-linkers (76), which demonstrated that increasing the cross-linker density markedly enhances the elastic stiffness while the frequency-dependent power-law scaling of the viscoelastic response remains largely unchanged. However, at longer timescales (> ∼ 10 s), we found faster stress relaxation for *Plec*^−/−^ cells than for *Plec*^+/+^ cells. At these long times, we expect stress relaxation to be tuned by the active turnover of the cytoskeletal filaments and/or rearrangements of the cytoskeleton via crosslinker exchange or myosin contractility (71). Interestingly, FRAP experiments on actin-GFP transfected cells demonstrated that actin turnover was faster in plectin knockout cells and coincided with the time scale of stress relaxation in step strain experiments. Taken together, this suggests that plectin changes viscoelastic relaxation mainly by impacting actin turnover. Intriguingly, a similar increase in actin turnover has also been observed in vimentin knockout cells, where the absence of vimentin led to enhanced phosphorylation of GEF-H1 at Ser886, thereby elevating RhoA activity and promoting stress fiber assembly, which in turn accelerates actin dynamics (77). However, we note that the environmental conditions in those experiments, where cells were adherent to flat substrates, are different from our experiments, where the cells were confined between nonadhesive plates. Consistent with these nonadherent conditions, confocal imaging did not reveal clear signs of stress fibers and only showed a fine actin meshwork. In future it will be interesting to investigate the dependence of actin turnover on plectin by comparing fibroblasts under different environmental conditions (2D versus 3D, adherent versus nonadherent).

Finally, our strain ramp experiments confirmed that plectin-deficient cells were softer than wild type cells at small deformations. However, at larger strains, both cell types showed stress-stiffening, characterized by a marked increase of the differential stiffness beyond a critical deformation. Notably, at large deformations, not previously accessed in other studies of the mechanical impact of plectin, plectin knockout cells reached a similar stiffness as wild type cells. The modulus-strain curves converged at high stress and showed a similar scaling with stress, suggesting that the mechanism of nonlinearity is the same. Actin and vimentin filaments are both semiflexible polymers. Theoretical models predict such semiflexible polymer networks to strain-stiffen due to the entropic cost of pulling out thermal bending fluctuations (78, 79). For uniform dense networks, models predict an affine response with modulus increasing with stress as *σ*^3/2^, consistent with experiments on reconstituted actin and intermediate filament networks (80, 81). In our experiments the stiffening response tended to a weaker scaling, with an exponent between 0.5 and 1. This could potentially signify a nonaffine response. The comparable stress-stiffening response of wild type and plectin knockout cells revealed by the collapse of our data onto a master curve suggests that the processes underlying nonlinear elasticity remain largely unaffected by plectin’s absence.

Strain ramps with repeated loading-unloading cycles revealed another interesting effect of plectin at large deformations. Wild type cells that had not yet been compressed had a higher linear modulus than plectin knockout cells, but after 1 or more additional cycles, the wild type cells became about as soft as plectin knockout cells. By contrast, the plectin knockout cells did not soften significantly during repeated loading. This finding suggests that plectin crosslinkers initially contribute to cell stiffness in wild type cells, but are disrupted by large compressive deformations. This suggests a model where plectin contributes an important mechanism of mechanical protection of the cytoskeleton (Figure 7). Upon uniaxial compression, due to its incompressible nature, the cell bulges at the edges, leading to stretching of the cytoskeleton (Figure 7A). In an unperturbed cell, actin and vimentin networks interpenetrate and are crosslinked by plectin (Figure 7B). With increasing deformations, stretching of the cytoskeleton first leads to the disruption of plectin crosslinking. We are not aware of any measurements of the plectin binding and unbinding rates to the cytoskeleton in cells. However, measurements on a biochemically reconstituted system showed that the PRD domains of plectin bind quite strongly to vimentin filaments, with a dissociation constant, *k*_d_, of −0.9× 10^−7^M (82), while the actin-binding domain binds more weakly to actin filaments (*k*_d_ = 3.2 ×10^−7^M) (83). Even if plectin could exhibit catch bond behavior like various actin-binding proteins (13), at sufficiently large mechanical load we expect plectin bonds to dissociate. With further increasing compressive strain, most likely the actin network is disrupted first. The more mechanically resilient vimentin network can likely remain intact, as indicated by measurements on reconstituted networks and on cells showing that intermediate filaments can sustain large (> 200%) strains (84).

**Figure 7:**
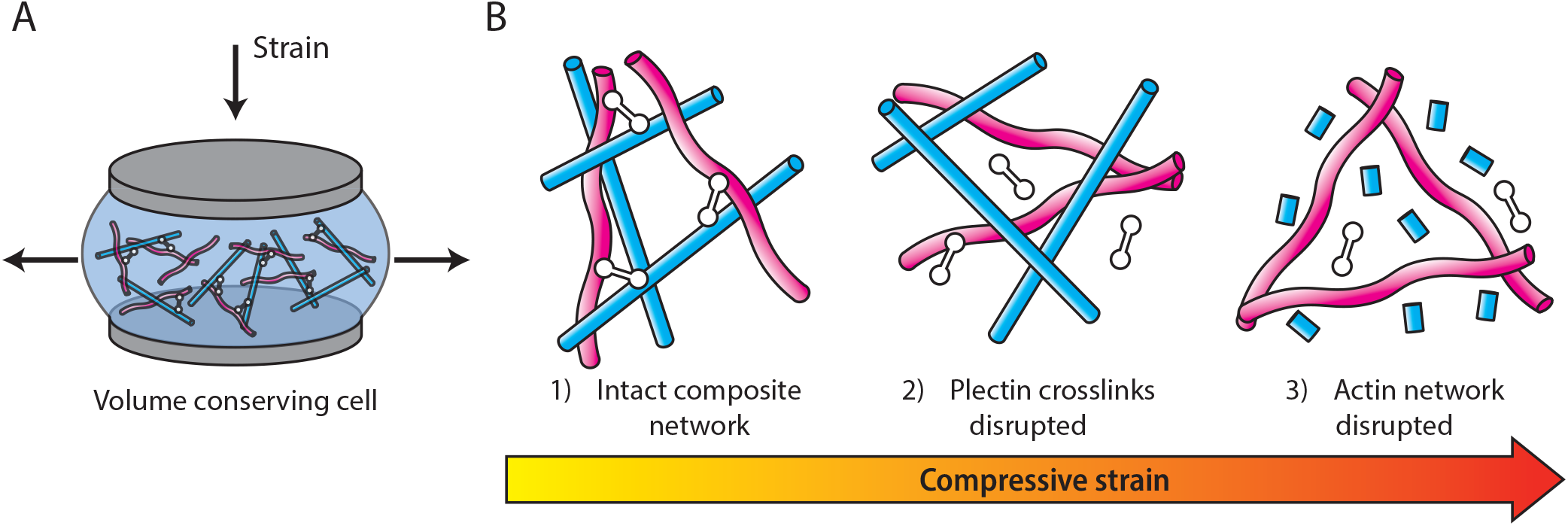
Proposed conceptual model for the impact of plectin on the strain-dependent mechanical response of single cells under compression. A) Upon uniaxial compression, due to its incompressible nature, the cell bulges at the edges leading to stretching of the cytoskeleton. (B) Starting with an intact composite actin-vimentin network crosslinked via plectin (left), stretching of the cytoskeleton first leads to the disruption of plectin crosslinking (middle). With further increasing compressive strain, the actin network is disrupted, leaving only the more mechanically resilient vimentin network intact (right).

Our results suggest that plectin contributes primarily to the elastic reinforcement of the cytoskeletal network at small deformations, while having minimal effect on the power-law rheology and stress-stiffening of fibroblasts. Crosslinking via plectin, mainly thought to be between actin and vimentin (19), makes a significant contribution to the rheology of fibroblasts.

However, plectin’s multifaceted roles complicate the interpretation of its mechanical contributions. In addition to crosslinking actin and intermediate filaments, plectin is known to interact with the linker of nucleoskeleton and cytoskeleton (LINC) complex (85, 86)—thereby linking the nucleus to the cytoskeleton—and to localize at focal adhesions where it contributes to adhesion dynamics and force transmission (35, 64). Consequently, the observed reduction in stiffness upon plectin deficiency may result not only from the loss of its crosslinking activity but also from disrupted connections at the nuclear envelope and altered focal adhesion mechanics. Future studies employing domain-specific manipulations will be essential to disentangle these distinct functions and to clarify the relative contributions of plectin’s various binding interactions to cellular mechanics.

## CONCLUSION

In this study, we have shown that plectin knockout (*Plec*^−/−^) fibroblasts are twice as soft as their wild type (*Plec*^+/+^) counterparts at small and large deformations, but their strain-stiffening response to compression is similar. Applying a step strain led to poroelastic stress relaxation at short times and a non-exponential response at long times with faster stress relaxation for *Plec*^−/−^ cells as compared to *Plec*^+/+^ cells. We hypothesize that this difference is due to a 3-fold difference in the kinetics of F-actin network turnover in the absence of plectin-mediated crosslinking to vimentin. When cells were subjected to repeated compression cycles, wild-type fibroblasts progressively softened to stiffness levels comparable to those of plectin knockout cells, indicating that plectin provides a first line of defense against large compressive loads. Confocal fluorescence images of the cells revealed an altered vimentin cytoskeletal network for *Plec*^−/−^ cells, where vimentin intermediate filaments formed bundles, rather than the finer meshwork displayed in *Plec*^+/+^ cells.

In conclusion, our findings reveal that plectin crosslinking significantly affects the cytoskeletal organization and viscoelastic properties of fibroblasts, at least partially through crosslinking of the actin and vimentin cytoskeletal networks. These findings provide a starting point towards understanding the role of mechanical integration of the different cytoskeletal networks in the cell interior and emphasize that these networks cannot be regarded as separate entities.

## AUTHOR CONTRIBUTIONS

J.P.C., M.G.L, F.C.M. and G.H.K. designed the research. J.P.C., M.G.L. and N.vV. performed research. J.P.C., M.G.L, F.C.M. and G.H.K. wrote the manuscript. L.W. and G.W. provided the cellular model systems and contributed to data interpretation. All authors edited the manuscript.

## ACKNOWLEDGMENTS

We thank M. Mavrakis (Institut Fresnel) for the vimentin-GFP plasmid used for live cell imaging experiments, C. Silva Martins (Institut Fresnel) for help with electroporation and G. Charras (University College London) for the actin-GFP plasmid used for FRAP measurements. We thank F. Ramirez Gomez for help with flow cytometry measurements and C. Sharma for helping with FRAP experiments. We thank A. Bonfanti for valuable help and advice regarding the RHEOS Julia package. G.H.K. gratefully acknowledges funding from the NWO Talent Programme, which is financed by the Dutch Research Council (project number VI.C.182.004). L.W. was funded by Austrian Science Fund (FWF) grants P31541-B27 (grant-DOI 10.55776/P31541) and I6049-B (grant-DOI 10.55776/I6049). F.C.M. was supported in part by the National Science Foundation Division of Materials Research (Grant No. DMR-2224030) and the Center for Theoretical Biological Physics (Grant No. PHY-2019745).

## SUPPLEMENTARY MATERIAL

An online supplement to this article can be found online

