## Supplementary figures, tables and methods for "Plectin affects cell viscoelasticity at small and large deformations"

### 1 Supplementary figures

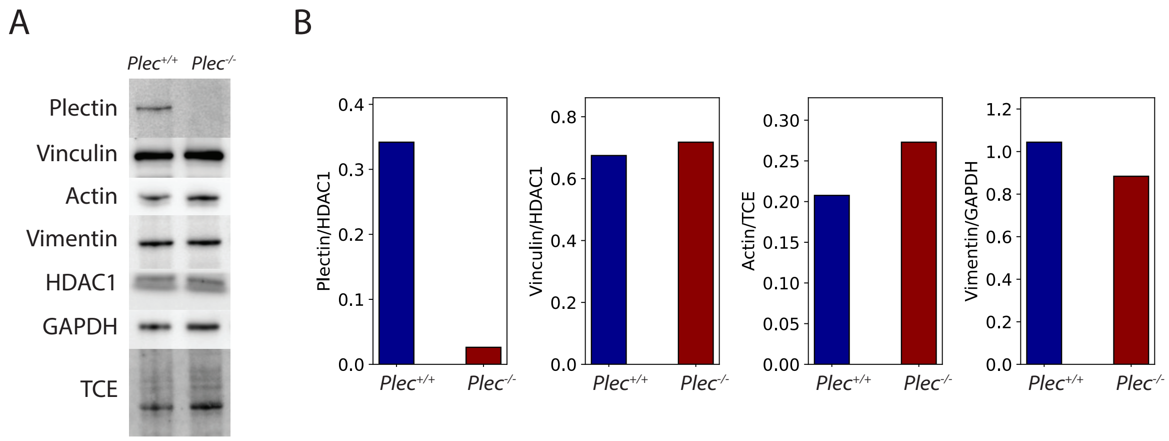


Figure 1: A) Protein expression levels of plectin, vinculin, actin and vimentin in Plectin+/+ and Plectin-/- mouse embryonic fibroblasts on cropped immunoblots. Depending on the protein of interest, HDAC1, GAPDH or TCE were used as a loading control. B) Quantification of the normalized band intensities as shown in panel A. Data shown for N=1 sample.

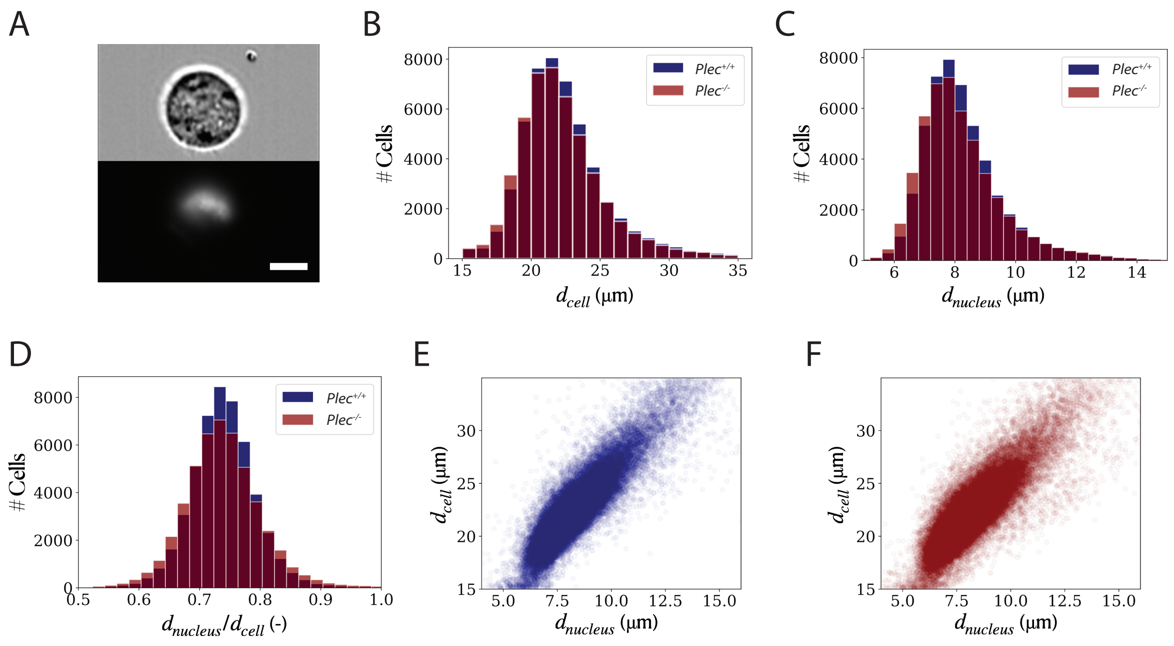


Figure 2: Cell and nucleus sizes measured by imaging flow cytometry for N = 50,220 *Plec*$^{+/+}$ cells and N = 48,711 *Plec*$^{-/-}$ cells. (A) Typical unprocessed bright field image of a cell (top) and corresponding epifluorescence image of its nucleus (bottom), which was stained with DRAQ5. Scale bar is 10 µm. (B) The mean cell diameters $d_{\text{cell}}$ are $22.4 \pm 3.1$ µm for *Plec*$^{+/+}$ cells and $22.2 \pm 3.1$ µm for *Plec*$^{-/-}$ cells. (C) Nuclear diameters $d_{\text{nucleus}}$ are $8.3 \pm 1.4$ µm for *Plec*$^{+/+}$ cells and $8.2 \pm 1.4$ µm for *Plec*$^{-/-}$ cells. (D) Nucleus-to-cell diameter ratio, $d_{\text{nucleus}}/d_{\text{cell}}$. On average, the ratio is the same for *Plec*$^{+/+}$ and *Plec*$^{-/-}$ cells ($0.74 \pm 0.06$ and $0.74 \pm 0.07$, respectively). Values stated as mean $\pm$ standard deviation. (E)-(F) Each symbol represents the $d_{\text{cell}}$ and $d_{\text{nucleus}}$ of a single cell. Both *Plec*$^{+/+}$ cells (E) and *Plec*$^{-/-}$ (F) cells display a proportional increase in size with their nucleus size.

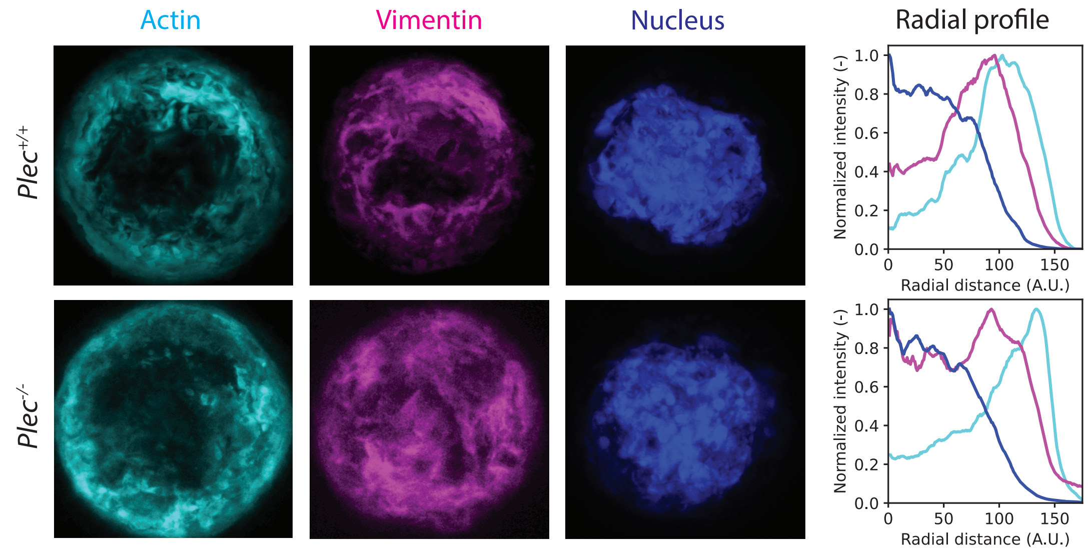


Figure 3: Actin, vimentin and nuclear organization in ‘average’ $Plec^{+/+}$ cells and $Plec^{-/-}$ cells. As we rescale cells to generate ‘average’ cells, scale bar not applicable. The images represent averages of a 2 µm thick confocal section at the cell equator for 10 $Plec^{+/+}$ and 10 $Plec^{-/-}$ cells. For both conditions, filamentous actin is located largely at the cell periphery, while vimentin is located closer to the center of the cell. Average radial profiles of the fluorescent signals (right) were calculated by taking normalized intensities around concentric circles from the center of the image, which show a significant overlapping region between actin and vimentin. The radial distance is set to zero at the cell center; the distance is in arbitrary units (see Methods for details).

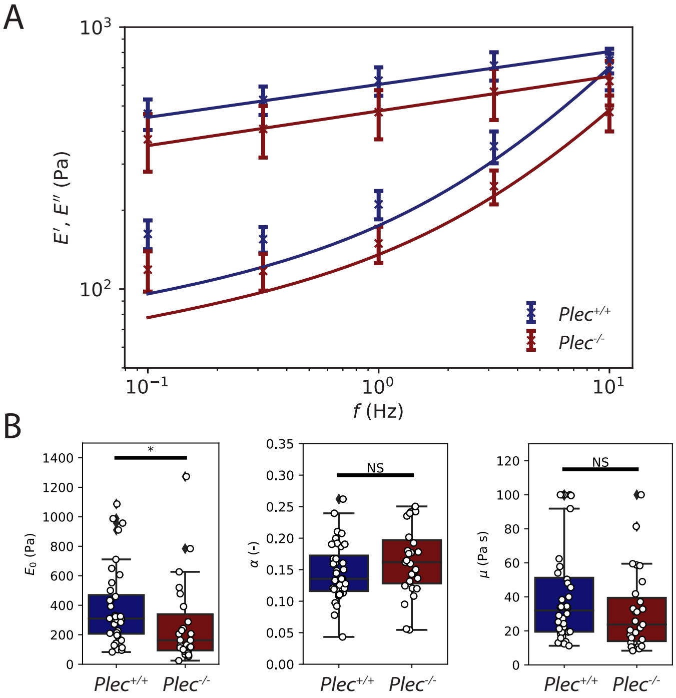


Figure 4: Small amplitude oscillatory strain measurements on single fibroblasts with fits to the structural damping model. (A) Storage ($E'$) and loss ($E''$) linear compressive moduli (symbols), measured for oscillation frequencies between 0.1Hz and 10Hz for $Plec^{+/+}$ (N=36) and $Plec^{-/-}$ (N=27) cells. Fits of the average data (solid lines) to the structural damping model (equations 1 and 2). The model does a poor job of capturing the mechanical behaviour of these cells at low frequencies. (B) We fit the rheological model to the individual cells to obtain mechanical fingerprints, i.e., scaling factor $E_0$, power law exponent $\alpha$ and cytoplasmic viscosity $\mu$. Asterisks indicate statistically significant differences (* denotes $P < 0.05$). N.S. denotes non-significant differences.

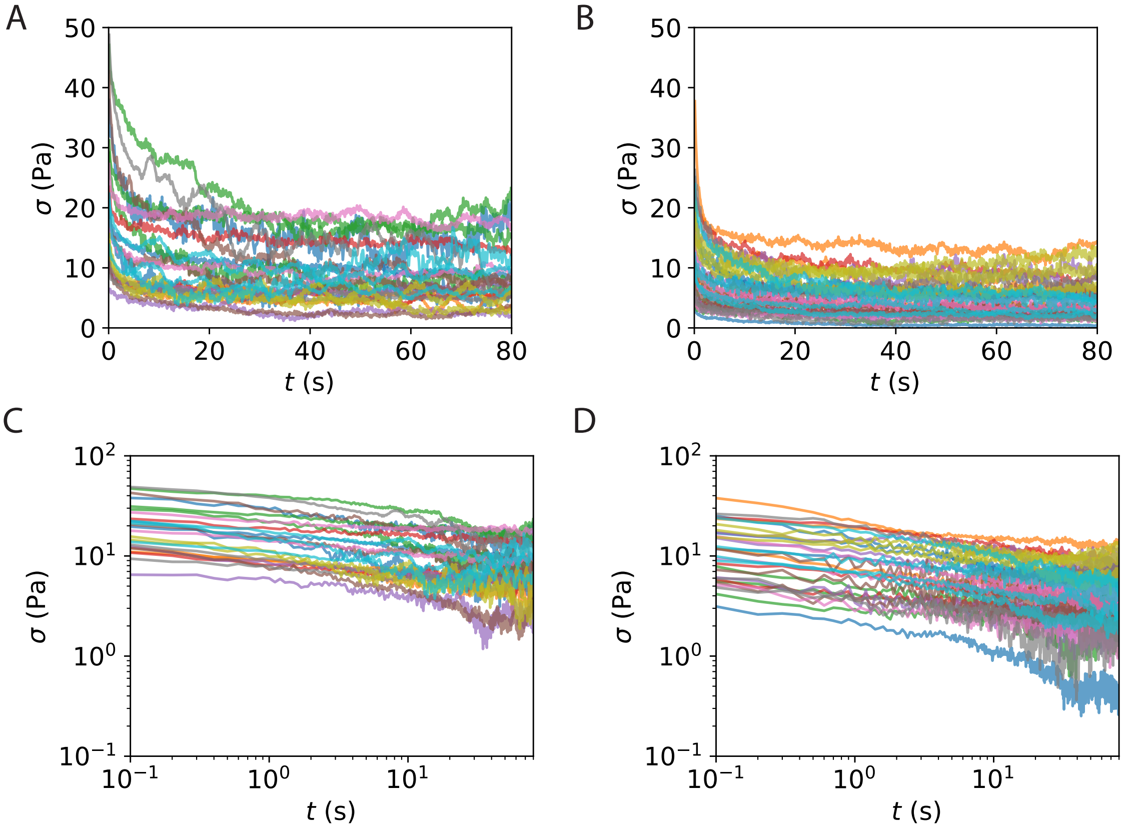


Figure 5: Individual stress relaxation measurements of the mechanical response of $Plec^{+/+}$ and $Plec^{-/-}$ cells to step strain compressions. We rapidly apply a compressive strain and hold it at a fixed value of 0.05. Subsequently, we monitor stress relaxation over 80 s. (A) Stress relaxation for all $Plec^{+/+}$ cells (N=25). (B) Stress relaxation for all $Plec^{-/-}$ cells (N=33). (C) We re-plot (A) on a log-log scale, as shown for averages in the main text. (D) We re-plot (B) on a log-log scale, as shown for averages in the main text.

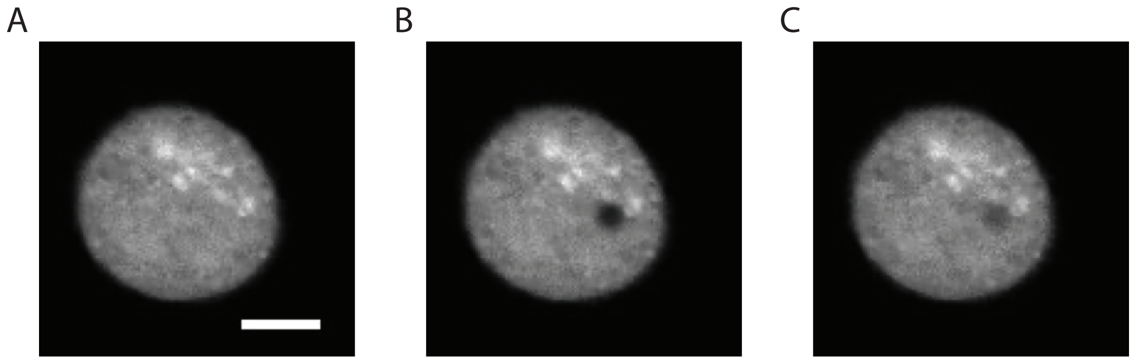


Figure 6: Fluorescence recovery after photobleaching (FRAP) on the cytoplasmic dye CellTracker Orange for a wild type fibroblast confined between two nonadhesive surfaces. The figure shows a representative example of (A) the pre-bleached cell, (B) the cell immediately after bleaching and (C) partial recovery of the fluorescence signal. Images are shown for a wild type cell. The FRAP experiment was performed in the mid-plane of the cell. Scale bar is 10 μm.

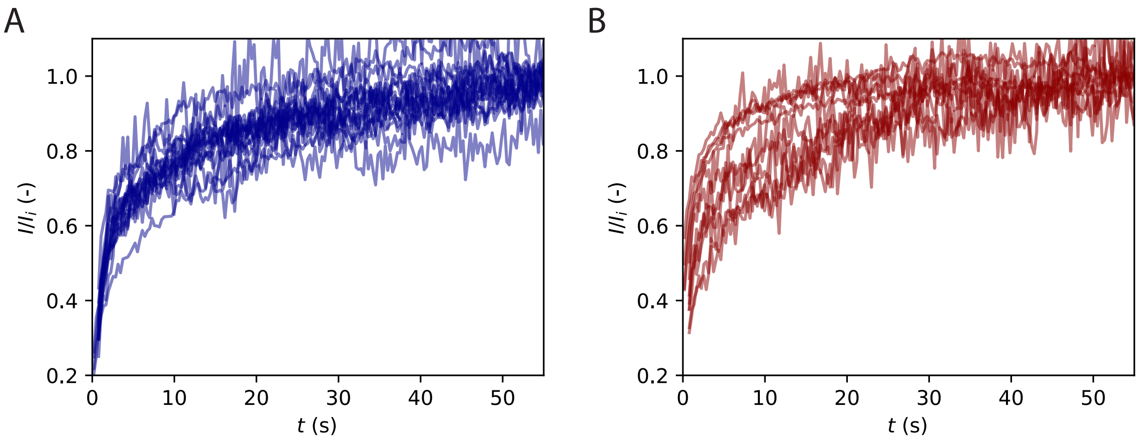


Figure 7: Individual fluorescence recovery curves for the comparison of actin turnover rates for $Plec^{+/+}$ cells (N=54) and $Plec^{-/-}$ cells (N=35) probed by fluorescence recovery after photobleaching (FRAP). (A) Individual normalized intensity curves for $Plec^{+/+}$ cells. (B) Individual curves for $Plec^{-/-}$ cells.

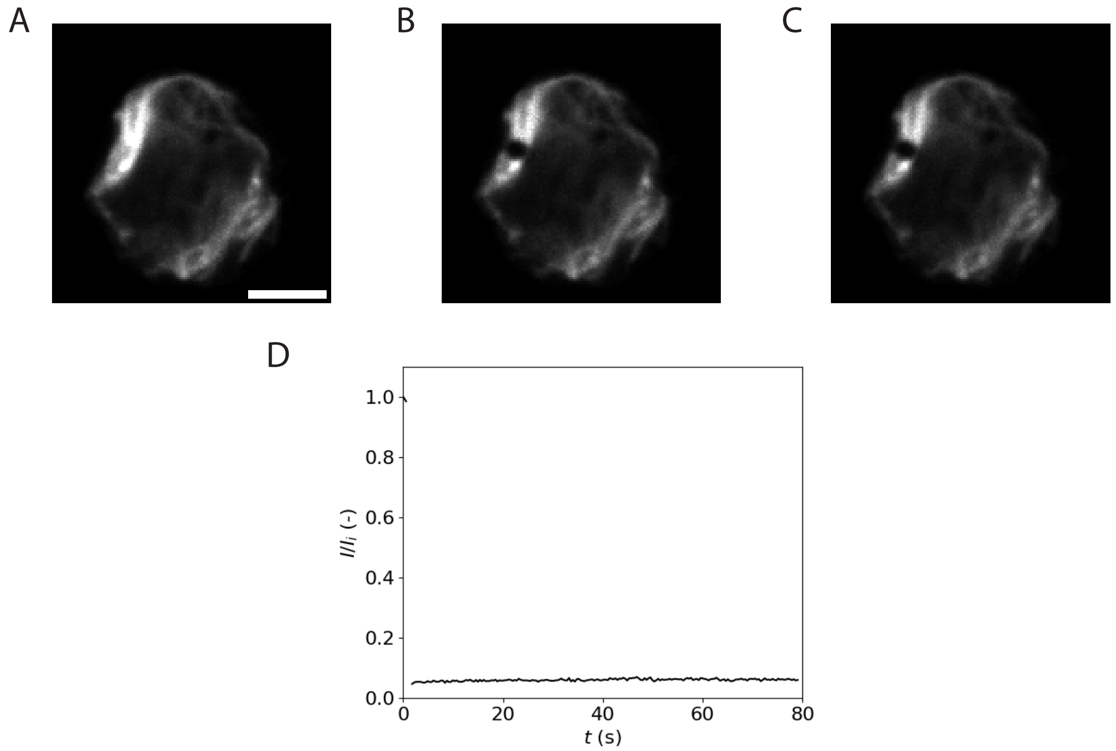


Figure 8: Fluorescence recovery after photobleaching (FRAP) experiments on vimentin-GFP tagged cells. Representative images of a FRAP experiment on a $Plec^{+/+}$ cell (A) before photobleaching, (B) immediately after bleaching, and (C) after 80 s of recovery. The FRAP experiment was performed in the mid-plane of the cell. Scale bar is 10 µm. (D) Representative recovery curve for the vimentin-GFP signal for a $Plec^{+/+}$ cell, showing negligible fluorescence recovery in 80 s (correspondin to the total duration of the stress relaxation experiments.

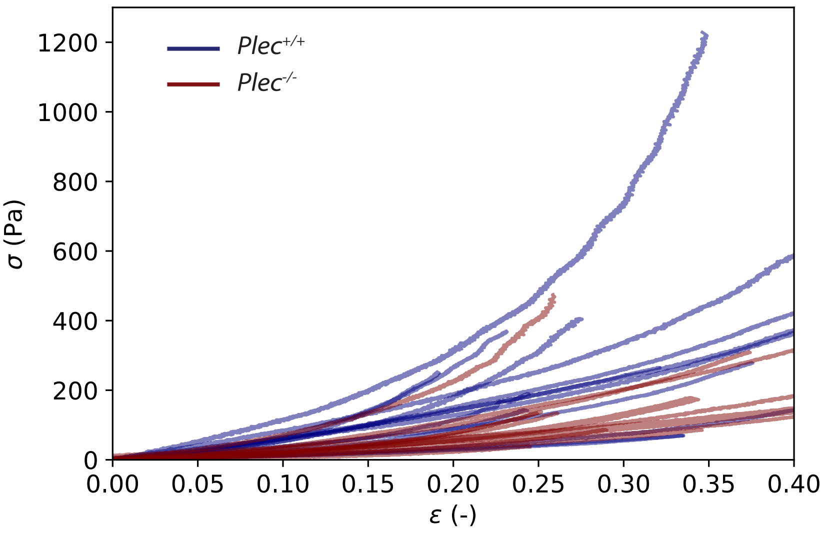


Figure 9: Individual strain ramp measurements of the nonlinear response of *Plec*$^{+/+}$ (blue, N=15) and *Plec*$^{-/-}$ (red, N=13) cells to large compressions, showing stress as a function of strain.

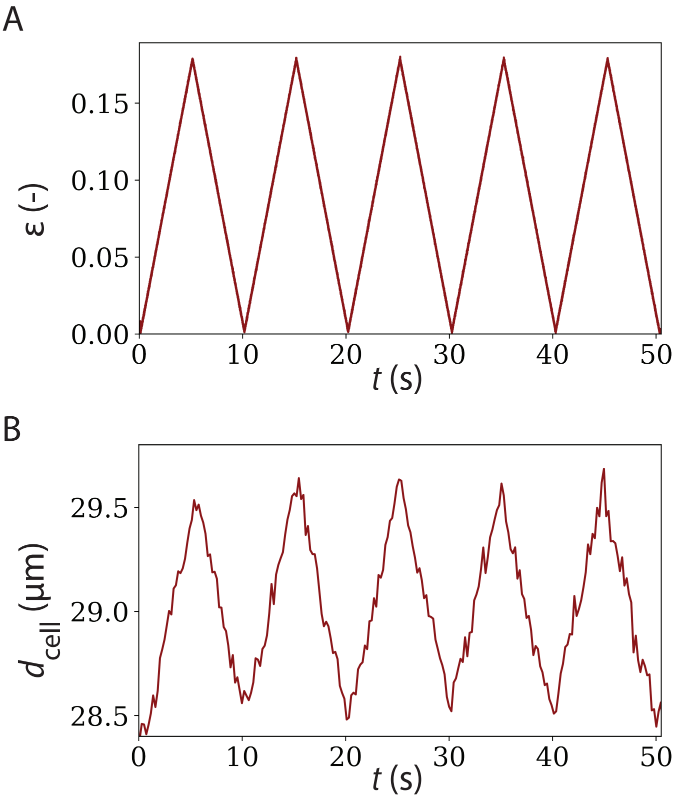


Figure 10: Representative cell diameter of a wild type fibroblast throughout 5 consecutive compressive loading-unloading cycles. (A) Strain profile during five consecutive strain cycles. (B) Measurement of the time-dependent changes in apparent cell diameter determined from the projected cell area observed by wide field epifluorescence imaging of CellTracker Orange. Changes in diameter are reversible.

#### 2 Supplementary tables

| **Primary antibody** | **Dilution** | **Manufacturer** | **Catalog number** |
|---|---|---|---|
| Rabbit anti-Plectin IgG | 1:1000 | Bioke | 12254s |
| Rabbit anti-Vinculin IgG | 1:500 | Thermofisher | 700062 |
| Mouse anti-Actin IgG | 1:500 | Millipore | MAB1501 Clone C4 |
| Mouse anti-Vimentin IgG1 | 1:2000 | Abcam | AB8978 |
| Rabbit anti-HDAC1 IgG | 1:1000 | ProteinTech | 10197-1-AP |
| Rabbit anti-GAPDH IgG | 1:5000 | Cell signal/Bioke | 2118s |

Table 1: Primary antibodies used for western blot analysis of protein expression levels of $Plec^{+/+}$ and $Plec^{-/-}$ cells.

| Secondary antibody | Dilution | Manufacturer | Catalog number |
|---|---|---|---|
| Rabbit anti-mouse IgG (HRP) | 1:5000 | Abcam | AB6728 |
| Goat anti-rabbit IgG (HRP) | 1:5000 | Abcam | AB97051 |

Table 2: Secondary antibodies used for western blot analysis of protein expression levels of $Plec^{+/+}$ and $Plec^{-/-}$ cells.

### 3 Supplementary methods

#### Fitting the structural damping law to oscillatory strain measurements

Before fitting the fractional viscoelastic model described in the main text, we first tried to capture the rheology of our cells using a simpler phenomenological model. This is a three parameter model referred to as the structural damping model [1–3], described in the main text as Equation 1. We take this equation and decompose it into its respective real and imaginary parts to yield the storage ($E'$) and loss ($E''$) moduli which we fit to our data:

$$E'(f) = E_0 c(\alpha) \cos\left(\frac{\pi\alpha}{2}\right)\left(\frac{f}{f_0}\right)^{\alpha} \tag{1}$$

$$E''(f) = E_0 c(\alpha) \sin\left(\frac{\pi\alpha}{2}\right)\left(\frac{f}{f_0}\right)^{\alpha} + \mu f. \tag{2}$$

The three fitting parameters are the scaling factor $E_0$, power law exponent $\alpha$, and cytoplasmic viscosity $\mu$. Using the curve fit function from Python's SciPy library [4], we simultaneously fitted Equations 1 and 2 to the measured storage and loss moduli.

#### Determination of cell and nucleus size by imaging flow cytometry

An ImageStreamX Mark II Imaging Flow Cytometer with a 40x objective was used to measure the sizes of the cells and their nuclei. One day before cytometry experiments, cells were detached with Trypsin-EDTA (0.25%) (25200056 Thermo Fisher Scientific) and counted using a Countess cell counter (Thermo Fisher Scientific). Next, 100,000 cells were transferred to plastic cell culture 6-well plates (83.3920.005 Sarstedt) containing culture medium. On the day of experiments, cell nuclei were fluorescently stained with DRAQ5 (P5739873 ThermoFisher) by adding DRAQ5 from a 5 mM stock solution in DMSO to the culture medium (1:500), and leaving the cells to incubate for 30 min. The cells were imaged in bright field, while the DRAQ5-stained nuclei were imaged in fluorescence with a 642 nm laser. The flow rate was set to Low in the ImageStreamX software, with High sensitivity, amounting to a flow speed of 55 mm/s. Single cells were separated from debris and cell clusters by selecting only objects with a diameter between 10 µm and 35 µm, a nuclear area of 50-5000 $\mu m^2$, and a cellular aspect ratio of 0.7-1. A typical image taken with the ImageStream flow cytometer is shown in Figure S2A. From the projected areas of the cells and nuclei, the corresponding average diameters were calculated, assuming spherical shapes in an automated Python script.

#### Monitoring the cell diameter during repeated ramp experiments

For experiments measuring the cell diameter during repeated compression, we used 35 mm glass bottom dishes (81218-200 Ibidi), in combination with a 40x NA 1.10 water immersion objective. To determine the cell diameter, we recorded each cell at a frame rate of 5fps. To determine the cell area, we thresholded each frame using Otsu's method in the Python library scikit image [5] and multiplied the number of pixels by the pixel size in the thresholded image. To determine the cell diameter, we used the geometric relation between a circle diameter and area: $A = \pi(d_{\mathrm{cell}}/2)^2$.

### Supplementary results

#### Flow cytometry

Complementing the images in the main text, we performed imaging flow cytometry measurements to determine cellular and nuclear sizes. The corresponding histograms are shown in Figure S2B-D. On average, *Plec*$^{-/-}$ cells have a diameter of $22.2 \pm 3.1$ µm, and are thus slightly smaller than *Plec*$^{+/+}$ cells with diameter $22.4 \pm 3.1$ µm. This diameter is calculated from the projected area $a$ as $d_{\mathrm{cell}} = 2\sqrt{a/\pi}$. In Figure S2B, we see that the diameters follow similar distributions, which appear to be log-normal distributions. Diameters range from 15 µm to 35 µm. The distributions for the nuclear sizes are shown in Figure S2C, where the diameter was again determined from the projected area. The resulting mean diameters $d_{\mathrm{nucleus}}$ are $8.3 \pm 1.4$ µm for *Plec*$^{+/+}$ cells and $8.2 \pm 1.4$ µm for *Plec*$^{-/-}$ cells. Dividing $d_{\mathrm{nucleus}}$ by $d_{\mathrm{cell}}$ for each cell yields the distribution shown in Figure 2D, with an average ratio of $0.74 \pm 0.06$ for *Plec*$^{+/+}$ cells and $0.74 \pm 0.07$ for *Plec*$^{-/-}$ cells. The ratios are thus comparable and both display a Gaussian distribution, with a slightly wider distribution for *Plec*$^{-/-}$ cells. This hints at a scaling between nucleus and cell size, which we confirmed by making scatter plots (Figure 2E-F).

#### Estimation of the poroelastic diffusion constant assuming fluid transport through the nucleus

In the main text, we make an estimate of the poroelastic diffusion constant, where we assume that the relevant length scale of fluid transport in the cell ($L$) is that of the cell membrane to the nucleus. If we instead use the cell size as the relevant length scale (i.e., assuming fluid transport occurs through cytoplasm and nucleus), we estimate significantly larger values for $D_{\mathrm{p}}$ of $(1700 \pm 100)$ µm$^2$/s for *Plec*$^{+/+}$ cells and $(3200 \pm 200)$ µm$^2$/s for *Plec*$^{-/-}$ cells. Importantly, we still observe a larger poroelastic diffusion constant in *Plec*$^{-/-}$ cells.

[4] Pauli Virtanen, Ralf Gommers, Travis E. Oliphant, Matt Haberland, Tyler Reddy, David Cournapeau, Evgeni Burovski, Pearu Peterson, Warren Weckesser, Jonathan Bright, Stéfan J. van der Walt, Matthew Brett, Joshua Wilson, K. Jarrod Millman, Nikolay Mayorov, Andrew R.J. Nelson, Eric Jones, Robert Kern, Eric Larson, C. J. Carey, İlhan Polat, Yu Feng, Eric W. Moore, Jake VanderPlas, Denis Laxalde, Josef Perktold, Robert Cimrman, Ian Henriksen, E. A. Quintero, Charles R. Harris, Anne M. Archibald, Antônio H. Ribeiro, Fabian Pedregosa, Paul van Mulbregt, Aditya Vijaykumar, Alessandro Pietro Bardelli, Alex Rothberg, Andreas Hilboll, Andreas Kloeckner, Anthony Scopatz, Antony Lee, Ariel Rokem, C. Nathan Woods, Chad Fulton, Charles Masson, Christian Häggström, Clark Fitzgerald, David A. Nicholson, David R. Hagen, Dmitrii V. Pasechnik, Emanuele Olivetti, Eric Martin, Eric Wieser, Fabrice Silva, Felix Lenders, Florian Wilhelm, G. Young, Gavin A. Price, Gert Ludwig Ingold, Gregory E. Allen, Gregory R. Lee, Hervé Audren, Irvin Probst, Jörg P. Dietrich, Jacob Silterra, James T. Webber, Janko Slavič, Joel Nothman, Johannes Buchner, Johannes Kulick, Johannes L. Schönberger, José Vinícius de Miranda Cardoso, Joscha Reimer, Joseph Harrington, Juan Luis Cano Rodríguez, Juan Nunez-Iglesias, Justin Kuczynski, Kevin Tritz, Martin Thoma, Matthew Newville, Matthias Kümmerer, Maximilian Bolingbroke, Michael Tartre, Mikhail Pak, Nathaniel J. Smith, Nikolai Nowaczyk, Nikolay Shebanov, Oleksandr Pavlyk, Per A. Brodtkorb, Perry Lee, Robert T. McGibbon, Roman Feldbauer, Sam Lewis, Sam Tygier, Scott Sievert, Sebastiano Vigna, Stefan Peterson, Surhud More, Tadeusz Pudlik, Takuya Oshima, Thomas J. Pingel, Thomas P. Robitaille, Thomas Spura, Thouis R. Jones, Tim Cera, Tim Leslie, Tiziano Zito, Tom Krauss, Utkarsh Upadhyay, Yaroslav O. Halchenko, and Yoshiki Vázquez-Baeza. SciPy 1.0: fundamental algorithms for scientific computing in Python. *Nature Methods*, 17(3):261–272, 2020.

[5] Stéfan Van Der Walt, Johannes L. Schönberger, Juan Nunez-Iglesias, François Boulogne, Joshua D. Warner, Neil Yager, Emmanuelle Gouillart, and Tony Yu. Scikit-image: Image processing in python. *PeerJ*, 2014(1):1–18, 2014.
